# Microbial diversity estimation and hill number calculation using the hierarchical Pitman-Yor process

**DOI:** 10.1101/2020.10.24.353599

**Authors:** Kevin Mcgregor, Aurélie Labbe, Celia M.T. Greenwood, Todd Parsons, Christopher Quince

## Abstract

**Background:** The human microbiome comprises the microorganisms that inhabit the various locales of the human body and plays a vital role in human health. The composition of a microbial population is often quantified through measures of species diversity, which summarize the number of species along with their relative abundances into a single value. In a microbiome sample there will certainly be species missing from the target population which will affect the diversity estimates.

**Methods:** We employ a model based on the hierarchical Pitman-Yor (HPY) process to model the species abundance distributions over multiple populations. The model parameters are estimated using a Gibbs sampler. We also derive estimates of species diversity, conditional and unconditional on the observed data, as a function of the HPY parameters Finally, we derive a general formula for the Hill numbers in the HPY context.

**Results:** We show that the Gibbs sampler for the HPY model performs well in simulations. We also show that the conditional estimates of diversity from the HPY model improve over naïve estimates when species are missing. Similarly the conditional HPY estimates tend to perform better than the naïve estimates especially when the number of individuals sampled from a population is small.

## 1. Introduction

Human microbiome studies attempt to quantify the makeup of the different species that occupy the human body. The composition of the gut microbiome in particular plays an important part in human health. Modelling species abundance distributions in this framework has long been a goal in metagenomic datasets. There have been a number of methods based on Bayesian non-parametrics developed to model these kinds of community dynamics. In particular, the Dependent Dirichlet process has been applied in this context to species abundance distribution and diversity estimation, and has additionally been used to address the effects of covariates on these entities (Ren et al., 2017; Arbel et al., 2016). However, these models were created from a statistical standpoint, and do not reflect the underlying principles of community ecology. In contrast, there are alternative statistical methods that attempt to describe and capture the dynamics of such distributions in a manner that is faithful to principles of ecological theory. Nevertheless, there are competing perspectives on how evolutionary processes and the environment affect species distributions (Jeraldo et al., 2012). For example, one theory suggests that species assemblage is characterized by niches, which are defined by the allocation of resources in the population (Chesson, 2000). There also exists a competing theory that describes “neutral” models, which assume that species are functionally equivalent and that processes such as immigration and birth/death events primarily contribute to community diversity (Etienne, 2005). The theory of neutrality claims that most genetic diversity observed in a population is a result of chance, as opposed to Darwinian selection (Kimura, 1968); that is, the birth rate that a particular species depends on the number of individuals of that species present in the population, rather than on the species’ ability to survive in the environment. This is analogous to the theory of neutrality for gene alleles that originated in the field of population genetics (Ewens, 1972). One important doctrine in neutral theory is Hubbell’s Unified Neutral Theory of Biodiversity (Hubbell, 2001). In Hubbell’s theory, it is assumed that there are a number of distinct local communities (sometimes called local populations) that are subject to immigration and birth/death processes. Immigration into each community is independent of the other communities, and all individuals that immigrate to a community do so from a conceptual *metacommunity* shared by all local communities. Immigration can happen at different rates across populations. The birth rate in the neutral model is on a *per capita* basis—births are more likely to occur for more highly abundant species. Additionally, the rate of speciation, i.e. the frequency at which new species appear (i.e. species that have not yet been observed in *any* population), is a defining parameter of the top-level metacommunity. Note that the metacommunity and the local communities in Hubbell’s model are sometimes referred to in the context of the *mainland-island* model, where immigration occurs from a mainland (analogous to metacommunity) to local islands surrounded by uninhabitable space (analagous to the local communities).

Harris et al. (2015) showed that the class of neutral models, when multiple local communities with differing population dynamics are considered and under certain conditions on individual mean reproductive success, converges to the hierarchical Dirichlet process (HDP) with large local population size. They developed a Gibbs sampler for the hierarchical Dirichlet process given a matrix of species counts among multiple local communities. The authors also presented a means for testing for neutrality in the populations after fitting the model. However, microbial abundance data has suggested that ranked species abundance distributions follow a power-law tail (Li, Bihan and Methé, 2013). That is, if we consider the ranked proportions of species in a population, *p*_*k*_, *k* = 1, 2, … such that *p*_1_ > *p*_2_ > *p*_3_ > …, then we have the following relationship for the ranked abundances:

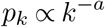

for some *a* > 0 (Clauset, Shalizi and Newman, 2009). One limitation of the Dirichlet process model is it cannot accommodate a power-law tail in its species distribution. Conversely, a related process called the *Pitman-Yor process*, can indeed generate a species distribution that exhibits a power-law tail (Goldwater, Johnson and Griffiths, 2006). It is then of value to pursue the question as to whether the Pitman-Yor model is more appropriate for modelling the species frequency distribution than the Dirichlet model. In this paper we investigate the use of the Pitman-Yor process in a hierarchical formulation called the *hierarchical Pitman-Yor process* (HPY process) to model abundance data in multiple microbial populations. The HPY model has been employed outside the context of community modelling (REFS). It has also been recently used in species discovery (Battiston, Favaro and Teh, 2018) as well as in single-cell RNA-Seq data (Camerlenghi et al., 2020). However, to the authors’ knowledge, this model has not previously been used in a species diversity estimation framework, nor have estimates of diversity metrics been derived in terms of the HPY model parameters. Therefore, in this article we provide such a derivation of diversity metrics in this context; in particular, we derive a general formula for the *Hill numbers* as a function of HPY parameters.

The HPY process is analagous to the mainland-island model, but it can also accommodate a departure from the assumption of neutrality. We develop a Gibbs sampler to fit the model parameters. In addition to the Gibbs sampler, we derive expressions for two measures of species diveristy in the context of the HPY process. We perform an extensive simulation study to investigate the performance of the HPY model under different situations, as well as the efficacy of the measures of species diversity defined in the HPY context.

We begin in the next section with the definition of the Pitman-Yor Process. In the Methods section (Section 2.1), we introduce the hierarchical Pitman-Yor model and give details for the Gibbs sampler as well as a description on calculating diversity in this model. In Section 3 we present results for an extensive simulation study that investigates the performance of the HPY model. Finally, in Section 4 we show results from applying the HPY model to a gut microbiome 16S sequencing dataset from a study on lean and obese twin pairs.

### 1.1. Pitman-Yor process

Assume that, for 0 *≤ α <* 1 and *γ* > *−α*, we generate a sequence of random variables *V*_*k*_, for *k* = 1, 2, … such that *V*_*k*_ *∼* Beta(1 *−α, γ* + *kα*). Furthermore, define the following:

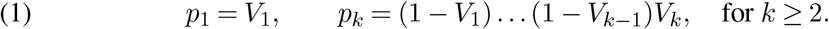

Then the *Griffiths, Engen, and McCloskey distribution* is defined as the joint distribution of (*p*_1_, *p*_2_, …) and is abbreviated as GEM(*γ, α*) (Yamato, Sibuya and Nomachi, 2001).

Let 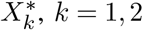,… be a sequence of independent samples from some distribution *H*. If we draw the vector *p* ∼ GEM(*γ, α*) independently from each 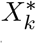, then the Pitman-Yor process (or two-parameter Poisson-Dirichlet process) is defined as:

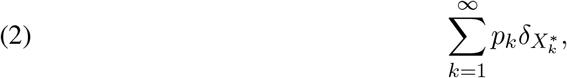

where 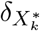 is a discrete measure at 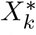 (Buntine and Hutter, 2010). The Pitman-Yor process is abbreviated as PY(*α, γ, H*). *α* and *γ* are called the *discount* and *concentration* parameters, respectively. *H* is referred to as the *base distribution*. The Dirichlet process is a special case of the Pitman-Yor process where the discount parameter *α* = 0.

A convenient way of drawing from the Pitman-Yor process is through the *Chinese restaurant process* representation. Assume that we have already drawn *X*_1_, …, *X*_*n*_ among which we have drawn the values 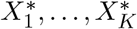 directly from the base distribution *H*, for some 1 *≤ K ≤ n*. If *H* is continuous, then all of 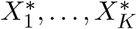 are distinct, however if *H* is discrete then some of the values may not be distinct. Then the distribution of *X*_*n*+1_ conditional on *X*_1_, …, *X*_*n*_ is:

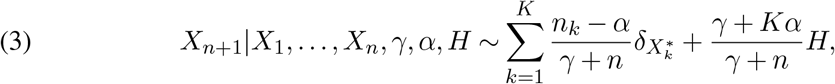

where *n*_*k*_ represents the number of times the value 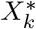 appears among *X*_1_, …, *X*_*n*_. More simply, *X*_*n*+1_ is sampled either from an existing value 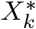 with probability proportional to *n*_*k*_, or from the base distribution *H*. The values 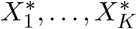 are called “tables” in the Chinese restaurant configuration. The tables may also be referred to simply by their indices 1, …, *K*.

### 1.2. Diversity

One of the most important descriptive tools in microbial community estimation is species diversity. Diversity aims to create a quantitative description of a population that incorporates information such as the total number of species in a population and the relative frequencies of those species. For example, one commonly applied measure of diversity is Simpson’s index, which concerns the probability of sampling the same species twice from a population. Assume a population contains *K* distinct species and that the proportion of species *k* is represented by *p*_*k*_. Simpson’s index is sometimes calculated as the probability of sampling two different species in subsequent draws (i.e. the complement of the above definition). The latter definition is used in this paper. Simpson’s index, denoted by *D*, is then written as:

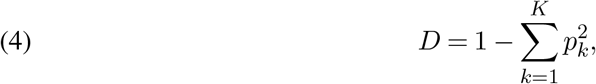

(Gorelick, 2006). A larger value of *D* implies the probability of obtaining the same species in two subsequent samples from the population is small. Thus, the closer the value of *D* is to one, the more diverse the population under consideration.

The fundamental problem in estimating diversity is that it is unlikely that a sample from a population will include all *K* species that exist in the population. To illustrate the problem, consider a population with three species (*K* = 3), where the counts of the three species are (*n*_1_, *n*_2_, *n*_3_) = (5, 4, 1). If we calculate Simpson’s index naïvely based on Equation 4 we get *D* = 0.58. Now, consider a sample from this population where we obtain the counts (*n*_1_, *n*_2_, *n*_3_) = (3, 2, 0). Calculating Simpson’s index on the sample proportions gives *D* = 0.48. Thus, the diversity in the population has been underestimated, simply because we failed to sample individuals from species 3.

Model assumptions can help to estimate species diversity in the presence of missing species. Methods for estimating Simpson’s index in the mainland-island model assumed in Hubbell’s neutral theory of biodiversity have been developed by Cerquetti (2015). Expressions to estimate Simpson’s index have not yet been developed in the HPY context, however. Thus, in this paper we propose the derivation of an expression of Simpson’s index in this context. The idea is that the HPY Gibbs sampler proposed in this paper could be applied to species abundance data, and estimates of the HPY model parameters could be obtained. From there, Simpson’s index could be calculated using the estimated HPY model parameters. The advantage of this approach is that, even if a particular population has missing species, the model structure assumed in the HPY framework can help to model the probability of discovering a new species (i.e. one that has not yet been observed). There is also a sense of “borrowing strength” between populations, in that we can account for sampling probabilities for a species that has a zero count in a population by considering the frequencies of that species in another population.

The framework that we pursue to estimate Simpson’s index using the HPY model depends on calculation of the *Hill numbers* (Hill, 1973), which are defined for integers *q ≥* 2 as:

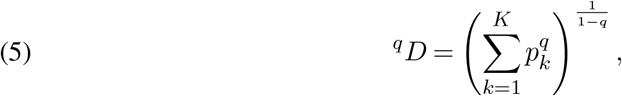

so that Simpson’s index as defined in Equation 4 can be written as

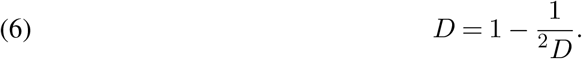

In this paper we show that the Chinese restaurant franchise representation of the HPY process admits a convenient way to derive the reciprocal of ^2^*D*. Additionally, we provide a formula for ^*q*^*D* for general *q ≥* 2 in the HPY context.

## 2. Methods

In this section we outline the model for the HPY process and describe the Gibbs sampler used to fit the model. Additionally we present a formula for Simpson’s index in the context of the HPY model parameters. In the mainland-island model, each individual in a local community can be traced back to an ancestor that immigrated to the community. In the hierarchical Pitman-Yor model, the “tables” in the Chinese restaurant representation of the Pitman-Yor process are analogous to individuals’ ancestors that immigrated in the mainland-island model. Multiple individuals in a community could have descended from the same ancestor and there can be multiple ancestors of the same species. The makeup of the ancestors are not directly observable and are therefore treated as latent variables in this framework. In this paper we use the following notation:

- *J* is the number of observed local communities
- *K* is the total number of observed species
- *Y*_*jk*_ is the observed count of species *k* in population *j*
- *t*_*jp*_ is the ancestor (“table”) from which individual *p* in community *j* descends
- *m*_*jk*_ is the number of ancestors in community *j* corresponding to species *k*
- *n*_*jtk*_ is the number of observations in community *j* that descended from ancestor *t*, corresponding to species *k*.
- *k*_*jp*_ is the species of individual *p* in community *j* (a number from 1 to *K*)
- *ψ*_*jt*_ is the species of the *t*^*th*^ ancestor in community *j* (a number from 1 to *K*)

We use the symbol *·* in a subscript to denote the summation of all values over a particular dimension. For example, 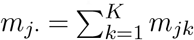 is the number of ancestors in population *j* over all species *k* = 1, …, *K*.

### 2.1. Hierarchical Pitman-Yor process model

In this section we introduce the hierarchical Pitman-Yor process and describe how it is connected to the mainland-island model. The model is defined as follows:

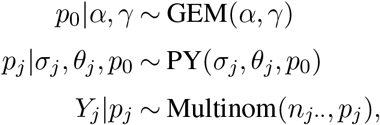

for *j* = 1, …, *J*. The vector of proportions *p*_0_ represents the abundance of species in the metacommunity which is assumed to follow a GEM distribution. Each vector *p*_*j*_ represents the abundance distribution of local population *j*, and is assumed to follow a Pitman-Yor process with base distribution *p*_0_. Therefore, sampling a new ancestor in local community *j* is akin to sampling from the abundance distribution *p*_0_ of the metacommunity. Finally, the vector of observed species counts in population *j*, denoted by *Y*_*j*_, is sampled from a multinomial distribution with its proportion vector set to *p*_*j*_. This final step represents the sampling procedure from the local community. The hierarchical Dirichlet process developed in Harris et al. (2015), was shown to be the large population size limiting distribution for Hubbell’s neutral model under certain conditions on individual mean reproductive success. In the hierarchical Pitman-Yor model, we include the discount parameters *α* and *σ*_*j*_; it has been shown that a non-zero value for the discount parameter corresponds to non-neutral assemblage (Crane et al., 2016). Thus, the discount parameter could be used as a measure of departure from neutrality. Figure 1 gives a schematic of the assumed HPY model assuming two local populations.

**Fig 1.**
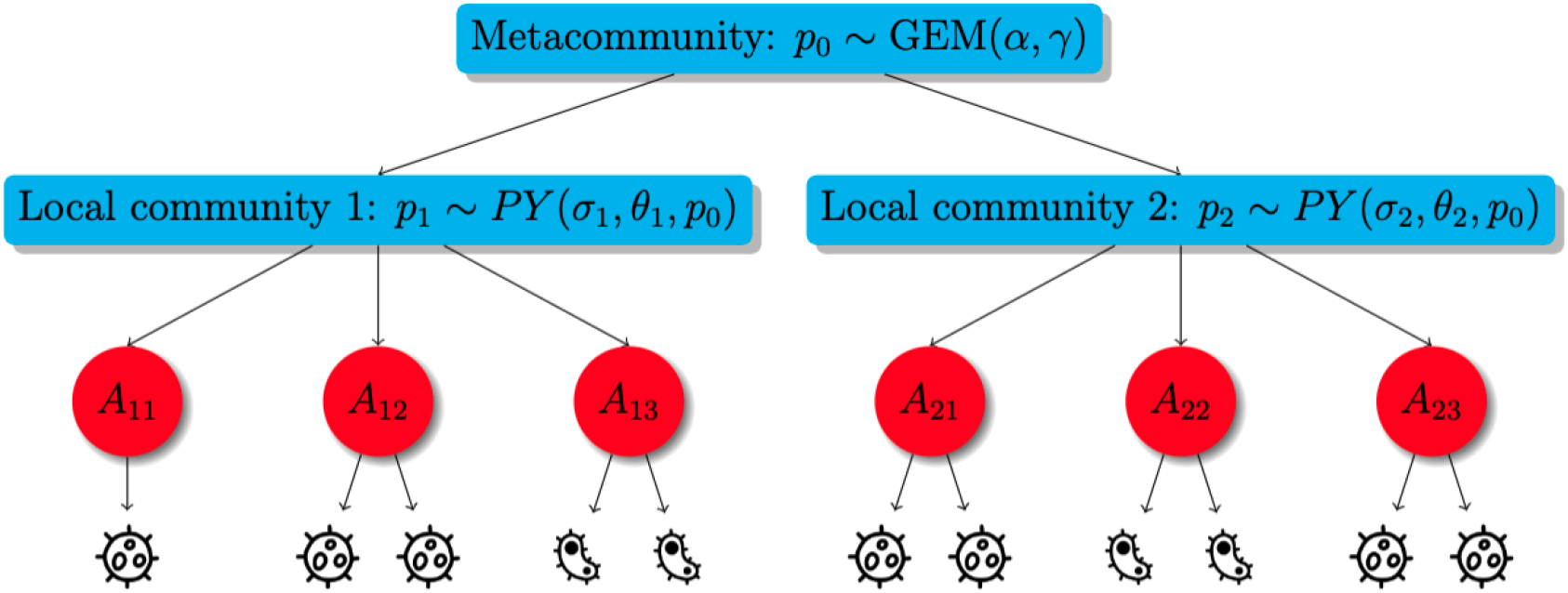
Schematic showing the HPY model assuming two local populations. A red node labelled Aij corresponds to ancestor j in local population i. The microbes pictured at the bottom correspond to descendants of the ancestors; i.e. the microbes present in the local communities. The observed species abundance table is a sample from the descendant microbes.

Priors for the top- and local-level discount and concentration parameters also must be specified in the HPY model:

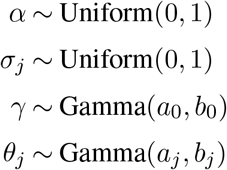

for *j* = 1, …, *J*. Here, *a*_0_ and *b*_0_ are respectively the shape and scale hyperparameters of the gamma distribution for the top-level concentration parameter *γ*. Likewise, *a*_*j*_ and *b*_*j*_ are the shape and scale hyperparameters in the prior for the local-level concentration parameter *θ*_*j*_.

### 2.2. Diversity estimation

In this paper we consider Simpson’s index in the context of the HPY model. The idea is, rather than calculating the Hill number reciprocal 1*/*(^2^*D*) directly from the observed proportions, we can infer in a given population the probability of sampling the same species twice in a row based on the sampling probabilities prescribed by the hierarchical Pitman-Yor process. This theoretical calculation would allow the inclusion of sampling species that have not yet been observed in a particular sample. We can do this by considering the Chinese restaurant process representation of the Pitman-Yor process. That is, when a particular species is sampled in a population, the probability of observing that same species a second time is slightly altered, since the corresponding ancestral counts will have been modified. Thus, the joint probability of sampling the same species twice can be calculated by considering all possible configurations of sampling the same species twice. We consider two forms of the expression for Simpson’s index: the first form, which we call the *unconditional* Simpson’s index, uses only the discount and concentration parameters from the HPY model; the second form, which we call the *conditional* Simpson’s index, uses the HPY parameters as well as the observed species counts and ancestral states *t*_*jp*_. The unconditional index represents the overall probability of sampling the same species twice in the HPY framework, and the conditional index represents the probability that the *next two* samples are of the same species, given a particular number of individuals have already been sampled. Considering the error in estimating the HPY discount and concentration parameters, we expect that additional leveraging of the observed counts and estimated ancestral states will aid in estimation accuracy for Simpson’s index. In addition to providing a means of calculating 1*/*(^2^*D*) in this way, we also provide a formula for calculating 1*/*(^*q*^*D*) for general *q ≥* 2; these results can be seen in Section S2.1.1.

Simpson’s index is a measure of alpha-diversity—in other words it measures the amount of diversity in a single population. In this paper we also consider the equivalent probability calculation across populations, i.e. the probability of sampling one species in one population and the same species in another population. This could be applied as a measure of beta-diversity, which measures the discrepancies in species abundance distributions between populations. This probability can similarly be calculated in the context of the hierarchical Pitman-Yor model. We have have derived expressions for Simpson’s index (conditional and unconditional) relating to both alpha-diversity and beta-diversity. As the expressions are quite complicated, they can be found in the supplement (Section S2).

### 2.3 Gibbs sampler for the HPY model

In order to fit the HPY model defined in Section 2.1, we use a Gibbs sampler. The parameters that must be updated in each step of the Gibbs sampler are the top-level GEM parameters *α* and *γ*; the local-level PY parameters *σ*_*j*_ and *θ*_*j*_, for *j* = 1, …, *J* ; and the ancestor indicators *t*_*jp*_ for each *j* = 1, …, *J* and *p* = 1, …, *n*_*j··*_.

As the local-level abundances are represented by a Pitman-Yor process, updates for the parameters *σ*_*j*_ and *θ*_*j*_ can be obtained by slice-sampling procedures defined by Buntine (2012). This algorithm is outlined in Algorithm S1 in the supplement. Note that the full conditionals for the concentration parameters are not log-concave. Buntine uses the technique from Escobar and West (1995) to generate an auxiliary beta-distributed random variable. The joint distribution of the concentration parameter and the auxiliary variable is log-concave (see Section S1.2.2) which allows the use of a slice-sampler. At the top level we have a GEM distribution, however the slice samplers from Buntine can still be applied with a slight modification to the table count vectors. Details on these procedures are given in Section S1.1 for the discount parameters, and Section S1.2 for the concentration parameters. For the ancestral indicators *t*_*jp*_, *j* = 1, …, *J* and *p* = 1, …, *n*_*j··*_, we apply the method used in Battiston, Favaro and Teh (2018). In their algorithm, each individual (i.e. each organism) is removed from the population and reassigned to a new or existing ancestor with probabilities defined by the Chinese restaurant process. Details of that procedure are outlined in Section S1.3. Algorithm 1 outlines a single iteration of the Gibbs sampler used to fit the model.

It is not immediately obvious what choices would be best for the hyperparameters *a*_0_, *b*_0_, *a*_*j*_, and *b*_*j*_. To guarantee the log-concavity of the joint distribution of the concentration parameters, we need to choose the shape parameters so that *a*_0_ *≥* 1 and each *a*_*j*_ *≥* 1 (see Section S1.2.2). We find that setting these shape parameters to 1.1 works well in practice. The sampler is fairly sensitive to the choice of the scale parameters *b*_0_ and *b*_*j*_. However, we do find in practice that setting all the scale parameters equal to the number of observed species *K* leads to good estimation accuracy. This allows the prior means for the concentration parameters to scale as a function of the number of observed species, which is necessary since we have observed that a poorly specified setting for this prior mean can negatively affect the posterior distributions of these parameters. This configuration corresponds to the recommendation in Buntine’s `libstb` C library, which is used to sample the Pitman-Yor parameters (Buntine, 2012).

### 2.4. Simulation study

To validate the Gibbs sampler for the HPY model, we have designed an extensive simulation study. We simulate data from the hierarchical Pitman-Yor process using its Chinese restaurant process representation. That is, for every new individual sampled in local population *j*, we will perform one of three options (1) sample from an existing ancestor, thus assigning that individual the same species as the ancestor; (2) sample from an existing species at the top level; or (3) sample a new species at the top level.

#### Algorithm 1

Iteration *i* of the HPY Sampler

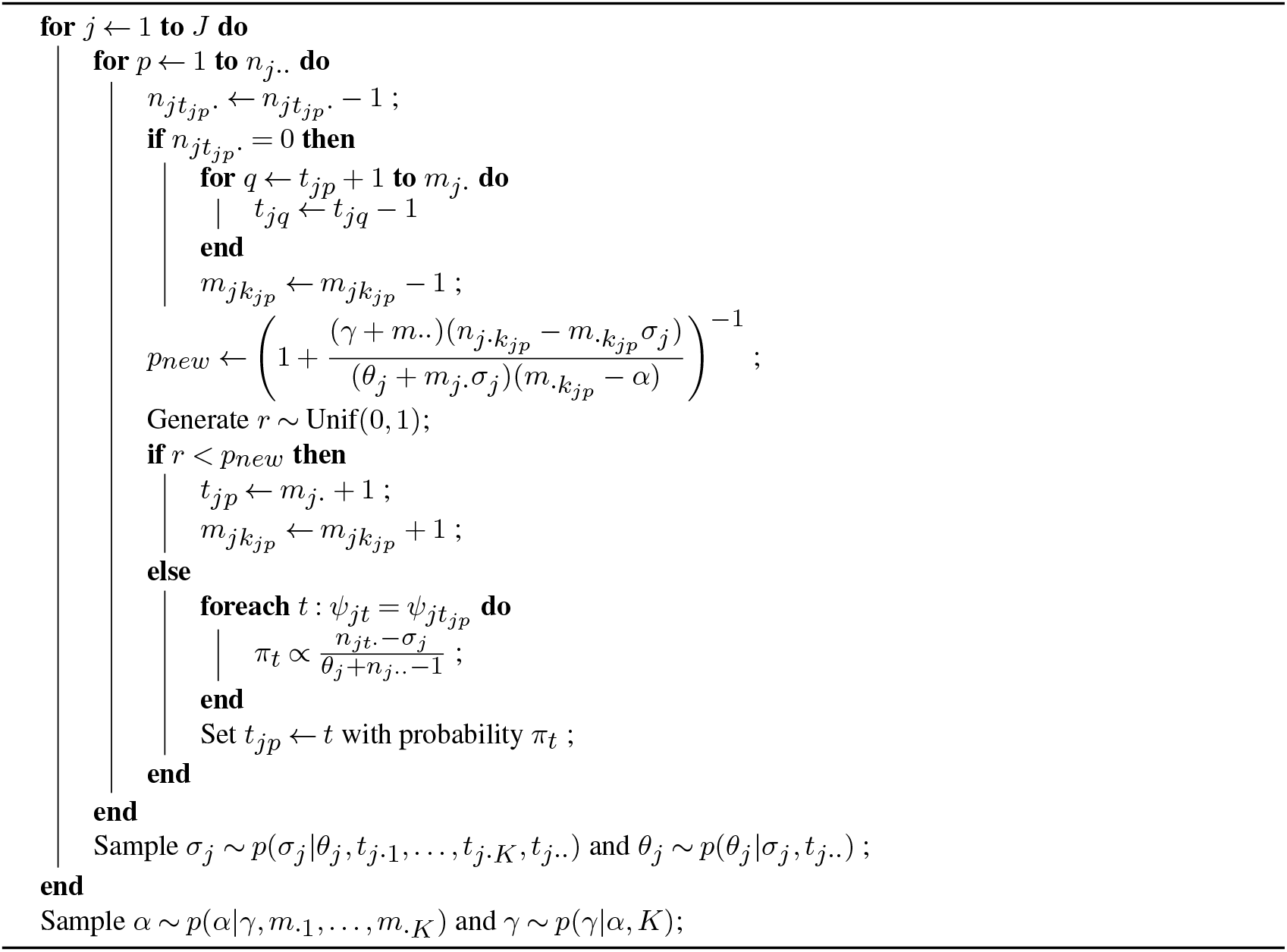

In the simulation study, we simulate data under multiple realistic configurations of the parameters in the HPY model. For the local-level parameters, we consider the values *σ*_*j*_ *∈* {0.2, 0.5} and *θ*_*j*_ *∈* {5, 25}. For the top-level parameters, we consider the values *α ∈* {0.2, 0.5, 0.8} and *γ ∈* {5, 25, 50}. We generate 10 local communities, and assume the same values for the local parameters across all communities. In all cases, the total number of individuals generated from the HPY process is 100,000 in each community; however, the final number of individuals sampled in each community in the multinomial step is *n*_*j··*_ *∈* {500, 1000, 5000} (i.e. this is the number of individuals that would be observed in the species abundance vector *Y*_*j*_ in each population *j*). We generated 50 replications of each simulation scenario. For each of the simulation replications the HPY Gibbs sampler was run for 2000 steps along with 1000 burn-in steps. To obtain parameter estimates we take the posterior mode of the MCMC samples. The posterior mode is more appropriate than the posterior mean in this context as the posterior distributions of the concentration and discount parameters are usually quite skewed due to the constraints on their supports.

Results from the simulation study are shown in Section 3. We examine the accuracy of the estimated parameters by comparing the posterior mode of the parameter samples from the Gibbs sampler to the true simulated values. We also calculate Simpson’s index to measure alpha-diversity in each population (see Equations S.13 and S.14) and beta-diversity to compare across populations (see Equations S.15 and S.16); we do this for both the unconditional and conditional versions of Simpson’s index. We compare Simpson’s index from the HPY model to the naïvely calculated indices, i.e. summing the squared observed proportions in each sample.

### 2.5. Data analysis in twin study

We apply the hierarchical Pitman-Yor model Gibbs sampler on a 16S sequencing dataset from the Missouri Adolescent Female Twin Study (Turnbaugh et al., 2009). The study was undertaken to investigate the differences in the composition of the gut microbiota between lean and obese hosts. The study considered monozygotic and dizygotic twin pairs that were concordant for obesity as well as their mothers, though we only include data from the twins in the HPY analysis. The study population consisted of 21-32 year-old women of European and African ancestry. 16S sequencing from fecal samples was performed using multiplex pyrosequencing on the V2 and V6 regions. Fecal samples were taken at two time points for each twin, however we consider samples only from the first time point, as there would likely be strong correlation between the compositions of the intestinal microbiota within each twin between the two time points. Most twin pairs lived apart, though 29% of the twin paris did live in the same household. In this analysis we consider a sample size of 102 samples from 29 monozygotic and 22 dizygotic twin pairs. In the cleaned dataset the are 28 women in the lean category (BMI 18.5–24.9 kg/m^2^), 7 women in the overweight category (BMI 25–30 kg/m^2^), and 70 women in the obese category (BMI > 30 kg/m^2^).

In this paper we wish to estimate diversity using the HPY model and compare diversity within the adiposity groups. We also explore some technical details such as the appropriateness of the fitted HPY model to the observed data and the effect of sequencing depth on the diversity estimates. From the HPY sampler we obtain 2000 MCMC samples after running 1000 burn-in samples. Note that, in consideration of the results of the simulation study, we only calculate the *conditional* version of Simpson’s index in this data application.

## 3. Simulation results

In this section we outline the results of the simulation study described in Section 2.4. First we consider the estimation accuracy of the various parameters in the HPY model (*γ, α, θ*_*j*_, *σ*_*j*_). Results from these investigations are shown in Section S3. Each of these plots shows estimation errors as the difference between the estimated parameter (posterior mode) and the true simulated value. These errors are normalized by dividing by the true simulated values to facilitate comparisons across simulation scenarios.

In Figure S1 we consider the normalized errors from estimating the top-level concentration parameter *γ*. Estimates generally appear to be unbiased. There is little improvement in estimation accuracy with increasing sample size in the local populations. Conversely, there does appear to be better estimation accuracy when the true value of the top-level discount parameter (*α*) is smaller. Higher values of *α* correspond to a heavier tail when considering the species abundance distribution aggregated across populations. That is, a higher top-level discount parameter implies many singleton species, which could contribute to the difficulties in estimation accuracy in this particular parameter configuration.

Simulation results for the top-level discount parameter *α* are shown in Figure S2. Again, the estimates do not appear to have any particular bias. In many cases there does appear to be a slight improvement in estimation accuracy with increasing sample size. There is a substantial increase in estimation accuracy for larger values of the true simulated value of *α*. In the *n*_*j··*_ = 500 configuration, there is generally better estimation accuracy for *σ*_*j*_ = 0.2 than for *σ*_*j*_ = 0.5.

In Figure S3 we show the estimation accuracy in simulations for the local-level concentration parameters *θ*_*j*_, *j ∈* {1, …, *J*}. There is generally better estimation accuracy when the true value of the local concentration parameters are *θ*_*j*_ = 5 compared to *θ*_*j*_ = 25. It also seems that there is a slight positive bias in estimates for this parameter, though the reason for this potential bias is still unclear. Similarly in Figure S4 we show results for the local-level concentration parameters *σ*_*j*_, *j ∈* {1, …, *J*}. In most cases there is a noticeable improvement with increasing sample size. In some cases there is an improvement for *θ* = 25 as opposed to *θ* = 5, however, this improvement is not consistent across all simulation scenarios. There also appears to be a slight negative bias for these parameters. In the MCMC samples there is typically a strong negative correlation between the local concentration and discount parameters due to the fact that both parameters are related to the total number of observed species. This could explain why the biases for the local concentrations are of the opposite sign of the biases of the local discount parameters.

Next, we investigate the results of estimating Simpson’s index using the HPY model. First, the unconditional and conditional versions of Simpson’s index are calculated from Equations S.13 and S.14, respectively. We calculate each index for each of the 2000 MCMC samples from the Gibbs sampler. Then, then final estimate for each version of Simpson’s index is taken as the mode of those values. It should be noted that the posterior distribution of Simpson’s index does not have a high variance, so the actual estimated value would not change much if the posterior mean or median were used. We compare the values of Simpson’s index from the HPY model to a naïvely calculated Simpson’s index from the observed proportions in each subsampled population. The true value of Simpson’s index is considered to be the result of applying Equation 4 to the proportions in the full simulated HPY dataset (which would be unobserved in practice).

In Figure 2 we compare the errors for the conditional and naïve estimation methods; the unconditional version also appears in Figure S5, but performs significantly worse than other methods. There is consistently greater variation in the amount of error for the unconditional version of Simpson’s index. For the sake of more easily making a comparison between unconditional and naïve, we have omitted the unconditional from Figure 2. The naïvely estimated Simpson’s index is often underestimated, especially when *γ* = 50, whereas the HPY conditional calculation was more accurate, though it did slightly overestimate in some cases, in particular when the sample size in the local populations was smaller. Recall that higher values of *γ* generally correspond to a higher species richness across all populations (i.e. in the metacommunity). Thus, the conditional Simpson’s index calculated using the HPY framework is more accurate when there is high species richness in the metacommunity. The conditional index is also generally more accurate for smaller population sizes. It should be noted that, as the number of samples taken within each population increases, both the naïvely calculated and HPY calculated values of Simpson’s index are consistently accurate. This shows that it is important to consider the number of sequences when calculating diversity in a population, and that the conditional estimates from the HPY model are valuable in samples with a low number of sequences.

**Fig 2.**
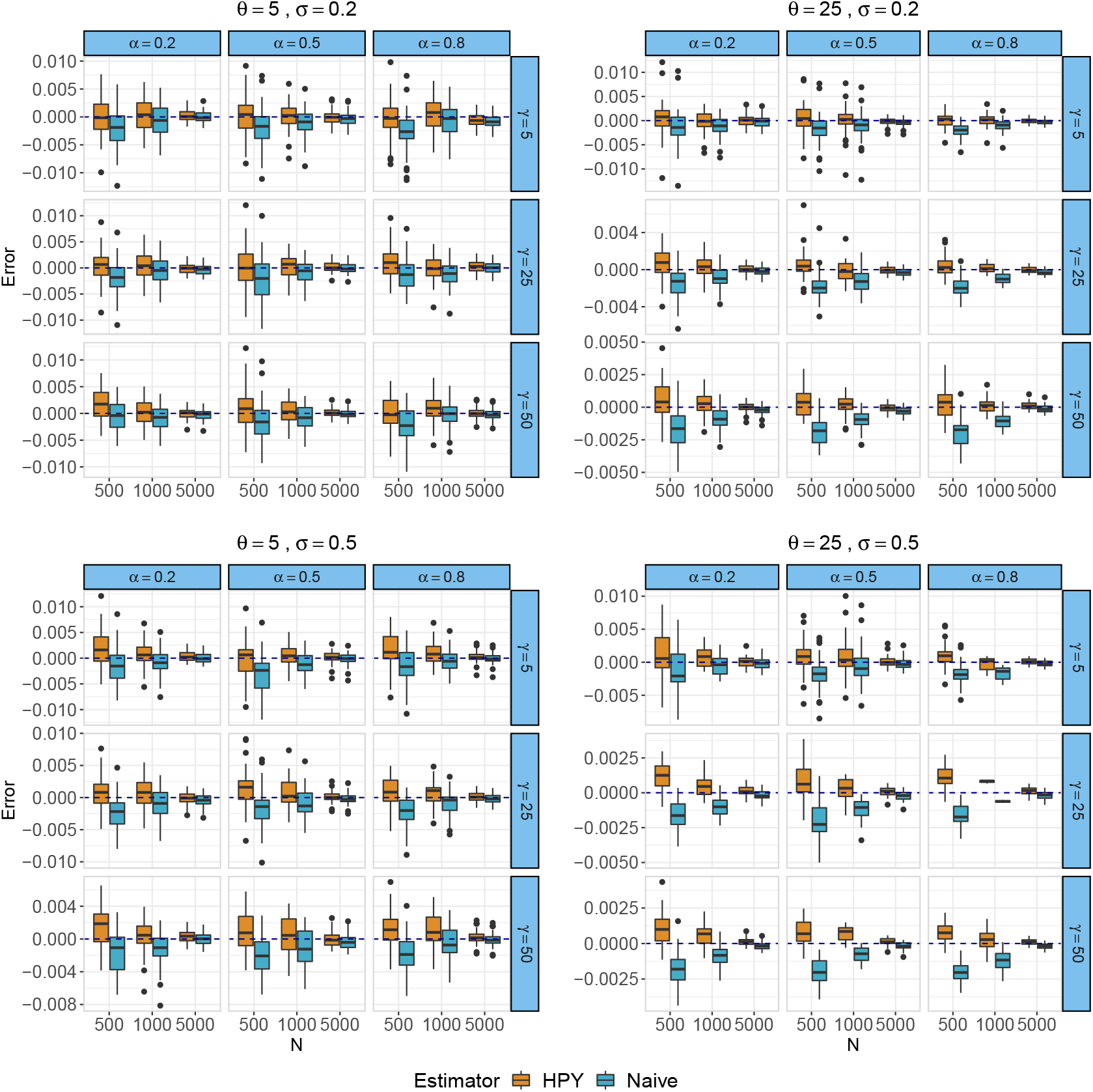
Estimation error for Simpson’s index comparing conditional Simpson’s index (HPY) and naïve estimates in simulated data.

We also explore the results of Simpson’s index for beta-diversity. In the HPY model we estimate this in the same way as for the alpha-diversity measure of Simpson’s index shown above, only we now use Equation S.15 for the unconditional index and Equation S.16 for the conditional index. Figure S6 shows the error for Simpson’s index for beta-diversity estimated from the HPY model (unconditional and conditional) and naïvely. There are not many discernible differences between the three estimation methods. This is likely due to the fact that, when calculating the underlying true beta-diversity values, it is probable that one or both of the proportions from a pair of populations will be zero. Thus, differences in the true beta-diversity values will be greatly affected by more abundant species, whereas differences in estimated values tend to occur due to differences in rarer species. Like before, estimation improves substantially for both methods with an increasing number of sequences.

## 4. Data analysis results

We now outline the results of applying the HPY model in the lean/obese twin study as described in Section 2.5. In this section we only consider the conditional HPY version of Simpson’s index, due to its clear advantage over the unconditional version as seen in the simulation study. First we consider the HPY estimated values of Simpson’s index within the obesity categories.

Figure 3(a) gives the distributions of Simpson’s index from the HPY sampler within each of the obesity categories. There is no apparent difference in alpha-diversity between subjects within the three categories using this measure. Similarly, we consider Simpson’s index for beta-diversity in Figure 3(b) calculated between subject pairs within the same obesity category and across obesity categories. Again, there is very little difference in beta-diversity using this metric in subject pairs concordant for obesity category compared to discordant.

**Fig 3.**
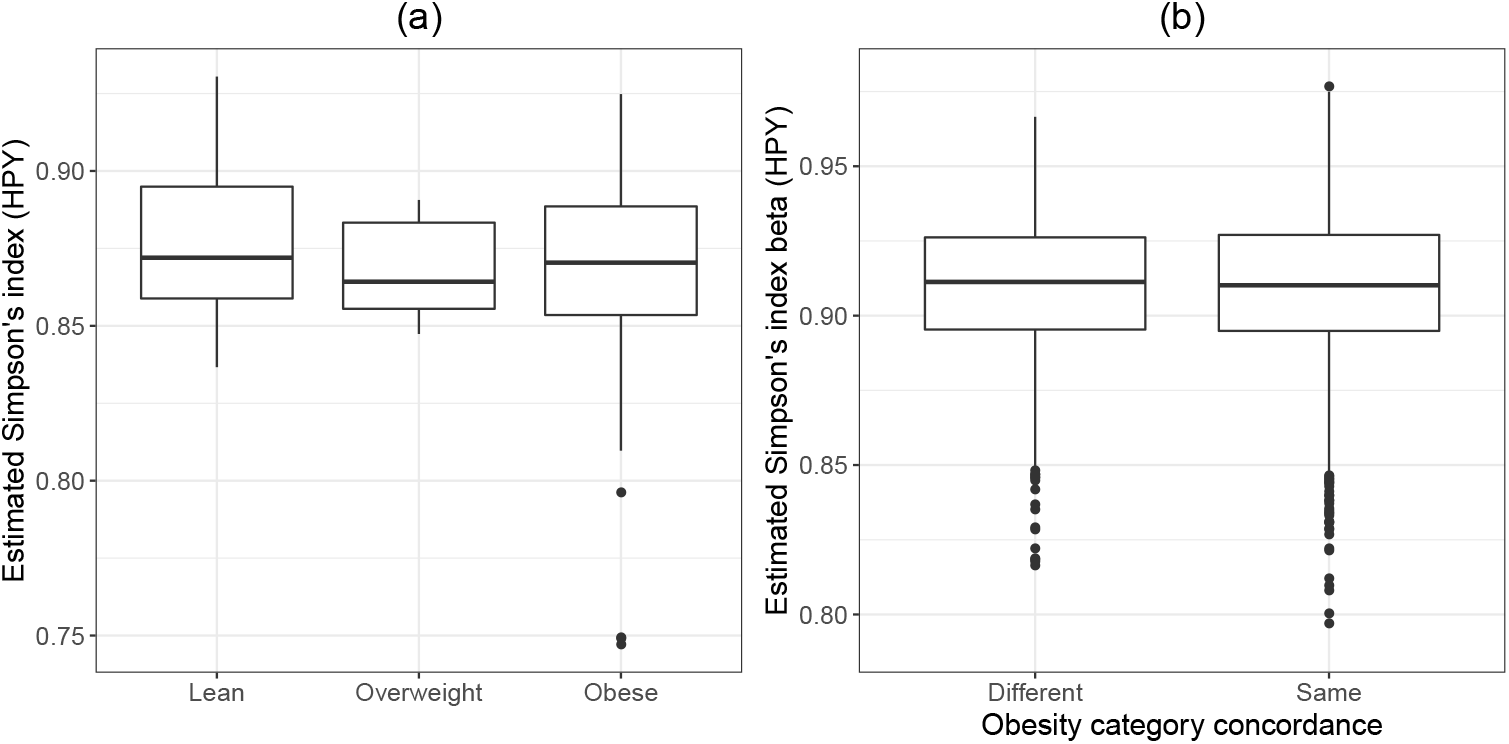
**(a)** Simpson’s index estimated from HPY model across overweight categories. **(b)** Simpson’s index for beta-diversity between subject pairs that are concordant vs. discordant for overweight categories.

We also check the distributions of the estimated values of the local-level HPY parameters in the obesity categories in Figure S7. As in the above results for diversity, there do not appear to be differences in the distributions of either the local concentrations *θ*_*j*_ or the local discounts *σ*_*j*_ between the obesity categories. Next, we check whether the estimated local-level parameter estimates correlate with the age of the host. Though there is no apparent relationship for *θ*_*j*_, there does appear to be a positive association between the age of the host and the local discount *σ*_*j*_, as shown in Figure S8. However, this association is rather weak with linear regression estimating a slope of only 0.0064, *p* = 0.026. Recall that a positive discount parameter reflects a non-neutral community. The range of estimated local discount parameters is from about 0.3 to 0.67, suggesting non-neutral assemblage within all the local communities.

Next we consider the effect of twin pairs on the results of the HPY model. In Figure S9 we examine scatterplots comparing (a) local concentration *θ*_*j*_, (b) local discount *σ*_*j*_, and (c) Simpson’s index between twin pairs. There is no apparent concordance for any of the three quantities between twin pairs. We also look at Simpson’s index for beta-diversity between and across twin pairs from the HPY model in Figure S10. There is a subtle difference in means of beta-diversity within and between twin pairs—the mean was 0.9098 for unrelated pairs, and 0.9027 within twin pairs. A two-sample t-test for comparing these means resulted in *p* = 0.0402. This means that the probability of sampling the same species twice is higher for twin pairs, suggesting slightly more similar diversity within twin pairs compared to unrelated pairs. Finally, we check whether alpha-diversity or beta-diversity differs with respect to monozygotic (MZ) or dizygotic (DZ) twin pairs in Figure S11. There does not appear to be any significant difference in either alpha or beta-diveristy between MZ and DZ twin pairs.

It is also important to check the appropriateness of the HPY model on this dataset. To do so, we compare the curve of ranked species abundances (averaged over all subjects in the dataset) to multiple datasets simulated under the HPY model. In each simulation we ran the HPY process using parameters from one of the MCMC samples from the Gibbs sampler run on the lean/obese twins dataset. We also did the same using a hierarchical Dirichlet process for comparison. Figure 4 shows the range of the the ranked species abundances curves for the HPY and Dirichlet models along with the observed species abundance curve from the dataset. It is immediately apparent that the HPY model fits the observed data much better than the hierarchical Dirichlet model. In particular, the HPY model better captures the tail of the distribution, which is unsurprising given the model’s ability to handle a power-law tail.

**Fig 4.**
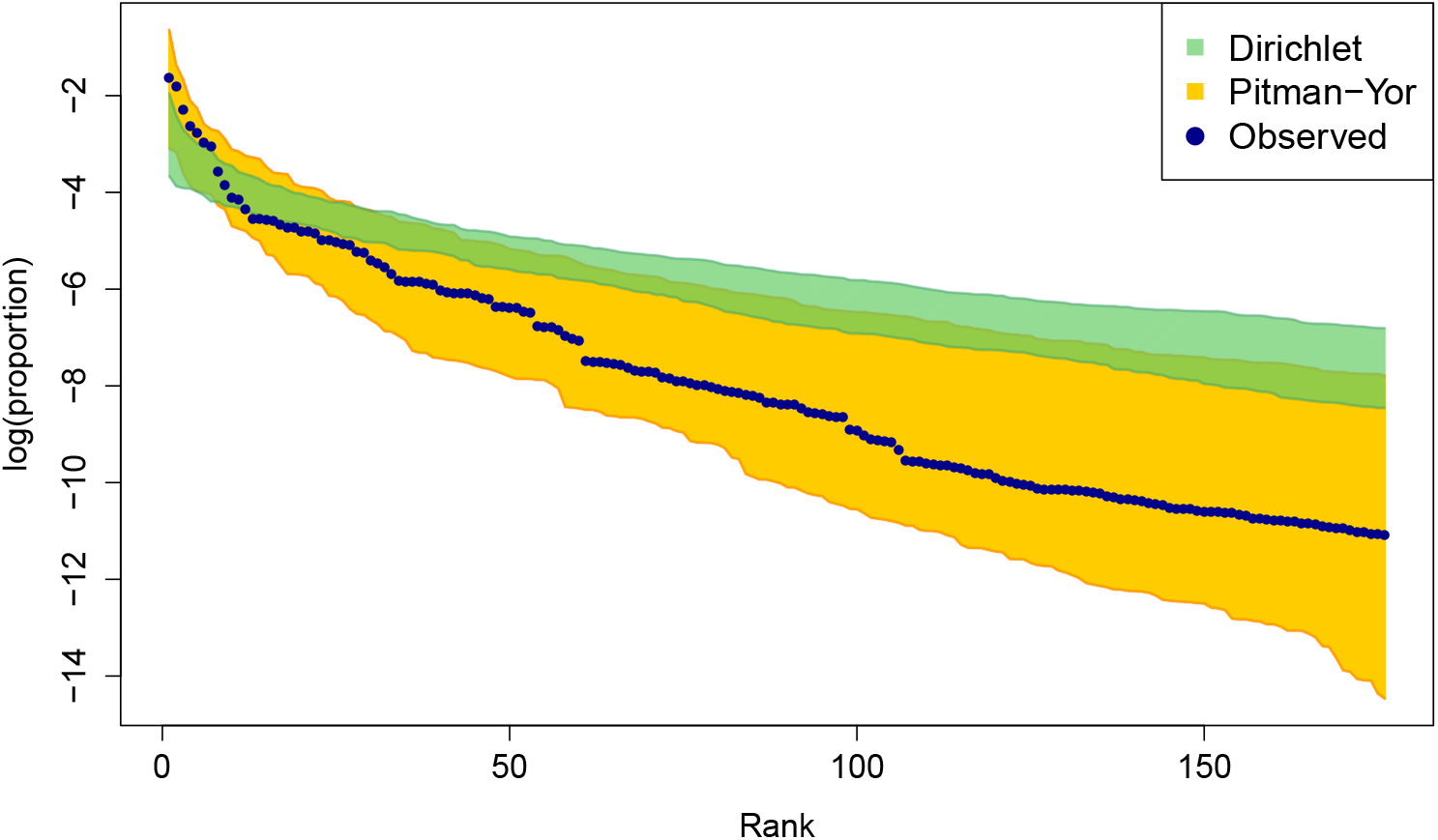
Ranked abundances of species in the twin study. The blue dots represent the true abundances and the shaded region represents the range of the ranked abundances in the simulated data from both the hierarchical Dirichlet and Pitman-Yor processes.

Finally we see the effect on the number of sequences in a sample on the difference in Simpson’s index (both alpha and beta-diversity) between the HPY model and naïvely estimated. In Figure S12 we see the differences in both indices as a function of the number of sequences; in the plot for beta-diversity we consider the sum of the sequences in both samples. In both cases there is more agreement between the HPY and naïve estimates with increasing sequence counts. This result is consistent with the simulation results presented in Section 3 and again underscores the fact that the HPY model is useful for estimating diversity in populations with a smaller number of sequences sampled.

## 5. Discussion

In this paper we have applied the hierarchical Pitman-Yor process to model species abundance distributions in microbiome sequencing data. We have developed a Gibbs sampler to fit the model and shown that the parameters in the HPY process are well estimated in this context. We have additionally provided a general formula for calculation of Hill numbers in the context of the HPY process. Finally, we have derived an estimator of Simpson’s index that leverages both HPY estimates from the Gibbs sampler as well as the observed data and inferred ancestral states. This conditional estimator was shown to outperform the unconditional estimator as well as the naïve estimator in most simulation cases.

The main limitation of the method is the computational inefficiency of the Gibbs sampler, in particular resampling the ancestral states *t*_*jp*_. Since sequencing depths could be quite high, there could be tens of thousands of ancestral states to update in each iteration of the Gibbs sampler. However, because all individuals of the same species within a population could be considered interchangeable to an extent, we do not have to keep track of the individuals’ ancestral states, rather we only need to keep track of the total number of ancestors corresponding to each species. In this case, ancestors can be added or subtracted from each population at a rate that depends on HPY parameters and current ancestor frequencies. Buntine and Hutter (2010) developed a sampler in this style which could be applied instead of the table indicator sampler as described by Battiston, Favaro and Teh (2018). Similarly, there have also been developments in variational approximations in Dirichlet mixtures that could be extended for the hierarchical Pitman-Yor model (Kurihara, Welling and Teh, 2007; Huynh, Phung and Venkatesh, 2016). Such approximations could greatly reduce the computational times in this model.

In this paper, when calculating Simpson’s index from the hierarhical Pitman-Yor model, we consider the results of sampling the *next two* individuals from the Chinese restaurant representation of the process, after having observed a sample from the target population. However this is distinct from the “true” diversity in a microbiome population, since the frequencies of the Chinese restaurant process will be different in the true underlying population. To get better estimates of diversity as well as species richness, individuals can be up-sampled within each population using the Chinese restaurant process. From there various measures of diversity could be calculated to estimate their true values in the underlying population. This could be useful in particular for diversity measures that would be difficult or impossible to write directly in terms of the HPY parameters.

One last point is that one of the main advantages of using the Pitman-Yor process instead of the Dirichlet process was to better capture the tail of the ranked species abundance distribution. However, since Simpson’s index is a diversity measure of second order, it is not sensitive to changes in the tail of this distribution. This means that Simpson’s index is not necessarily the best diversity measure for investigating the tail. However, the calculation of Simpson’s index can be expressed very nicely in terms of the probabilities defined by the Chinese restaurant representation. We believe using this index is a good starting point for demonstrating how diversity measures could be estimated using the HPY process, though future work should focus on alternative metrics to better capture the tail of the ranked species abundance distribution.

## 6. Conclusion

We have developed a Gibbs sampler to fit a model based on the hierarchical Pitman-Yor process given species abundance data across multiple microbial populations. This process is more appropriate for use on microbiota species abundance data than the previously established hierarchical Dirichlet process due to its ability to accommodate a power-law tail. Additionally, we have provided new expressions for Simpson’s index measuring alpha-diversity and beta-diversity in the context of the hierarchical Pitman-Yor process and showed their usefulness in simulation and in a data application. In doing so, we also derived a general formula for calculation of the Hill numbers in the HPY framework.

## Acknowledgements

K.M. would like to acknowledge support from an NSERC Discovery Grant (RGPIN-2021-03634). K.M. would like to acknowledge travel support from the Queen Elizabeth Scholars Program (“Quantitative Biology and Medical Genetics for the World”) as well as a McGill Graduate Mobility Award. K.M. was also supported by a Fonds de recherche du Québec-Santé doctoral award, as well as the McGill University Faculty of Medicine’s Cameron-Davis and Davis Fellowship and Gerald Clavet Fellowship. C.G. and A.L. acknowledge the CIHR operating grant (MOP-130344). C.G. would also like to acknowledge support from Compute Canada (ID 2541) as well as support from the Ludmer Centre for Neuroinformatics and Mental Health. C.Q. is funded through an MRC fellowship (MR/M50161X/1) as part of the Cloud Infrastructure for Microbial Genomics (CLIMB) consortium (MR/L015080/1). Images in Figure 1 used under CC BY 3.0 license. Attribution: microbe bacterium, bacillus, infusoria, microorgan by Alina Oleynik and Microbe by Sidiq Fathurochman from the Noun Project.

## SUPPLEMENTARY MATERIAL

### supplement.pdf

Supplementary file containing additional information relevant to the manuscript.

### R-package

Implementation of the Gibbs sampler and calculation of Simpson’s index in the HPY model. Package `hpy` is available at https://github.com/kevinmcgregor/hpy

### Twin study data

This dataset is publicly available: https://pubmed.ncbi.nlm.nih.gov/19043404/

