## Supplemental material for "Microbial diversity estimation and hill number calculation using the hierarchical Pitman-Yor process"

### Supplement: Microbial community modelling and diversity estimation using the hierarchical Pitman-Yor process

#### S1 Sampling the model parameters

##### S1.1 Sampling the discount parameters $\sigma_j$ and $\alpha$

The conditional distribution of the discount parameter $\sigma_j$ for local population $j$ can be expressed as:

$$p(\sigma_j|\theta_j, t_{j\cdot 1}, \ldots, t_{j\cdot K}, t_{j\cdot\cdot}) \overset{\sigma_j}{\propto} \sigma_j^{t_{j\cdot\cdot}} \frac{\Gamma(t_{j\cdot\cdot} + \theta_j/\sigma_j)}{\Gamma(\theta_j/\sigma_j)} \prod_{k=1}^{K} S_{t_{j\cdot k},\alpha}^{n_{j\cdot k}} \tag{S.1}$$

where $S_{M,a}^{N}$ is a generalized Stirling number of type $(-1, -a, 0)$, as defined by Hsu and Shiue (1998). Then, if we denote the log-probability density function for the full conditional of $\alpha$ by $\log(p(\alpha|\gamma))$, we can simply use the slice sampler as defined in Algorithm S1. Since $0 < \alpha < 1$, there is not the same concern about the initial value for the slice sampler as we have for the concentration parameter (see Section S1.2). An arbitrary starting value (say 0.5) can be used.

At the top level we have a GEM distribution, meaning that the full conditional distribution of the top-level discount parameter $\alpha$ can be written as (Buntine and Hutter, 2010):

$$p(\alpha|\gamma, m_{\cdot 1}, \ldots, m_{\cdot K}) = \alpha^K \frac{\Gamma(K + \gamma/\alpha)}{\Gamma(\gamma/\alpha)} \prod_{k=1}^{K} \frac{\Gamma(m_{\cdot k} - \alpha)}{\Gamma(1 - \alpha)} \tag{S.2}$$

We then exploit the following property for a special case of the generalized Stirling number $S^n_{1,a} = \Gamma(n - a)/\Gamma(1 - a)$. This allows Equation S.2 to be rewritten as.

$$p(\alpha|\gamma, m_{\cdot 1}, \ldots, m_{\cdot K}) = \alpha^K \frac{\Gamma(K + \gamma/\alpha)}{\Gamma(\gamma/\alpha)} \prod_{k=1}^{K} S^{m_{\cdot k}}_{1,\alpha}. \tag{S.3}$$

This form is equivalent to that of the full conditional for the discount parameter in the Pitman-Yor process with all the table counts fixed to 1. Thus we can modify Buntine's slice sampler to sample from the conditional distributions of top- and local-level concentration parameters. It has been shown that the full conditionals are log-concave (Buntine, 2012).

#### S1.2 Sampling the concentration parameters $\theta_j$ and $\gamma$

We wish to use a slice sampler to sample from the full conditional of the concentration parameters $\gamma$ and $\theta_j$, $j = 1, \ldots, J$. However, the full conditionals are not in fact log-concave. Instead we use the auxiliary variable method from Escobar and West (1995). For the concentration parameter for local population $j$, we can define an auxiliary beta-distributed random variable $p(q_j|\theta_j) \sim \text{Beta}(\theta_j, n_{j\cdot\cdot})$.

The joint distribution of $\theta_j$ and $q_j$ given $\sigma_j$ is written as:

$$\begin{aligned}
p(\theta_j, q_j|\sigma_j, t_{j\cdot\cdot}) &= p(\theta_j|\sigma_j, t_{j\cdot\cdot})p(q_j|\theta_j) \\
&\overset{\theta_j}{\propto} e^{-\theta_j/b}\theta_j^{a-1} \frac{\Gamma(t_{j\cdot\cdot} + \theta_j/\sigma_j)}{\Gamma(\theta_j/\sigma_j)} \frac{\Gamma(\theta_j)}{\Gamma(\theta_j + n_{j\cdot\cdot})} \frac{\Gamma(\theta_j + n_{j\cdot\cdot})}{\Gamma(\theta_j)\Gamma(n_{j\cdot\cdot})} q_j^{\theta_j - 1}(1 - q_j)^{n_{j\cdot\cdot} - 1} \\
&\overset{\theta_j}{\propto} e^{-\theta_j/b}\theta_j^{a-1} \frac{\Gamma(t_{j\cdot\cdot} + \theta_j/\sigma_j)}{\Gamma(\theta_j/\sigma_j)} q_j^{\theta_j - 1}(1 - q_j)^{n_{j\cdot\cdot} - 1}.
\end{aligned} \tag{S.4}$$

Now, because of the fact that:

$$p(\theta_j|\sigma_j) = \int_0^1 p(\theta_j|\sigma_j)p(q_j|\theta_j)dq_j, \tag{S.5}$$

we can first sample from the joint distribution of $\theta_j$ and $q_j$, and simply disregard the $q_j$ samples.

For the top-level concentration parameter, the full conditional is derived by considering the probability function of the GEM distribution. We can use the same auxiliary random variable procedure as we used for the local-level concentration parameter. That is, assuming $p(q_0|\gamma) \sim \text{Beta}(\gamma, m_{..})$ we have that:

$$p(\gamma, q_0|\alpha, K) \overset{\gamma}{\propto} e^{-\gamma/b_0}\gamma^{a_0-1}\frac{\Gamma(K+\gamma/\alpha)}{\Gamma(\gamma/\alpha)}q_0^{\gamma-1}(1-q_0)^{m_{..}-1} \tag{S.6}$$

In Section S1.2.2 we show that the probability function in Equation S.4 is log-concave in $\theta_j$, which allows the use of a slice sampler. However, because the support of $\theta_j$ is $(-\sigma_j, \infty)$, it can be difficult to choose an appropriate starting value for the slice sampler. To chose the starting value we use the fixed-point optimizer from Buntine (2012) to first find the maximum of $p(\theta_j, q_j|\sigma_j)$ and use that as the initial point of the slice sampler.

##### S1.2.1 Finding the maximum of $p(\theta_j, q_j|\sigma_j)$

The support of $\theta_j$ is $(-\sigma_j, \infty)$. Consequently, a poor choice of the initial value for the slice sampler could result in the sampler taking a very long time to warm up. Instead, we begin the sampler from a maximum a posteriori estimate of $p(\theta_j, q_j|\sigma_j)$. Differentiating the joint log-probability function of $\theta_j$ and $q_j$ gives:

$$\log(p(\theta_j, q_j|\sigma_j)) = -\frac{\theta_j}{b} + (a-1)\log(\theta_j) + \log\left(\Gamma(\theta_j/\sigma_j + t_{j\cdot\cdot})\right) - \log\left(\Gamma(\theta_j/\sigma_j)\right) \\ + (\theta_j - 1)\log(q_j) + (n-1) + c_{-\theta_j} \tag{S.7}$$

where $c_{-\theta_j}$ is a constant not depending on $\theta_j$. Setting $\frac{\partial \log(p(\theta_j, q_j|\sigma_j))}{\partial \theta_j}=0$ gives:

$$-\frac{1}{b} + \frac{a-1}{\theta_j} + \psi(\theta_j/\sigma_j + t_{j\cdot\cdot})\frac{1}{\sigma_j} - \psi(\theta_j/\sigma_j)\frac{1}{\sigma_j} + \log(q_j) = 0 \tag{S.8}$$

where $\psi(\cdot)$ is the digamma function. To solve this, Buntine (2012) designed the following fixed-point optimizer:

$$\theta_j^{(t)} \leftarrow \sigma_j \psi^{-1}\left[-\frac{\sigma_j}{b} + \frac{\sigma_j(a-1)}{\theta_j^{(t-1)}} + \psi\left(\frac{\theta_j^{(t-1)}}{\sigma_j} + t_{j\cdot\cdot}\right) + \sigma_j \log(q_j)\right] \tag{S.9}$$

where $\psi^{-1}(\cdot)$ is the inverse of the digamma function. This optimizer converges in a relatively small number of iterations (generally less than five). The output of this optimizer is then used as the initial value of the slice sampler for $p(\theta_j, q_j|\sigma_j)$.

##### S1.2.2 Log-concavity of $p(\theta_j, q_j|\sigma_j)$

The slice sampler requires $p(\theta_j, q_j|\sigma_j)$ to be log-concave. To show that this is indeed the case, we consider the second derivative of $\log(p(\theta_j, q_j|\sigma_j))$. If we differentiate left-hand side of Equation S.8 we get:

$$\frac{\partial^2 \log(p(\theta_j, q_j|\sigma_j))}{\partial \theta_j^2} = -\frac{a-1}{\theta_j^2} + \frac{1}{\sigma_j^2}\Big(\psi_1(\theta_j/\sigma_j + t_{j\cdot\cdot}) - \psi_1(\theta_j/\sigma_j)\Big) \tag{S.10}$$

where $\psi_1(\cdot)$ is the trigamma function (i.e. the second derivative of the logarithm of the gamma function). The trigamma function is strictly decreasing on the positive real line, meaning that:

$$\psi_1\left(\frac{\theta_j}{\sigma_j} + t_{j\cdot\cdot}\right) < \psi_1\left(\frac{\theta_j}{\sigma_j}\right) \tag{S.11}$$

since $t_{j\cdot\cdot} \geq 1$. Therefore, we have that $\frac{\partial^2 \log(p(\theta_j, q_j|\sigma_j))}{\partial\theta_j^2} < 0$ if $a \geq 1$.

#### S1.3 Sampling the ancestral states $t_{jp}$

Recall that each $t_{jp}$ indicates the ancestor (analogously called a "table" in the Chinese restaurant construction) from which the $p^{th}$ individual in population $j$ descends. As the true ancestral states of the individuals in the observed populations are unknown, this uncertainty must be accounted for in the model. Each ancestral state indicator $t_{jp}$ can be thought of as a random variable and can consequently be re-sampled in each iteration of the Gibbs sampler.

All individuals sharing a common ancestor must be of the same species. The species of each individual in the population is determined by the sample. Therefore, when considering the full conditional distribution of each $t_{jp}$, we must also condition on the observed species of individual $p$. Here we use the method presented in Battiston et al. (2018) wherein each ancestral state is updated by first "removing" individual $p$ from its population and either reallocating it to an existing ancestor (of the same species) or allocating a new ancestor. That is, we can sample $t_{jp}$ proportional to:

$$\sum_{t:\psi_{jt}=\psi_{jt_{jp}}} \frac{n'_{jt\cdot} - \sigma_j}{\theta_j + n_{j\cdot\cdot} - 1}\delta_t + \frac{\theta_j + m'_{j\cdot}\sigma_j}{\theta_j + n_{j\cdot\cdot} - 1}\frac{m'_{\cdot k_{jp}} - \alpha}{\gamma + m'_{\cdot\cdot}}\delta(m'_{j\cdot} + 1). \tag{S.12}$$

where $n'_{jt\cdot}$, $m'_{j\cdot}$, $m'_{\cdot k_{jp}}$, and $m'_{\cdot\cdot}$ are the revised values of $n_{jt\cdot}$, $m_{j\cdot}$, $m_{\cdot k_{jp}}$, and $m_{\cdot\cdot}$ after removing individual $p$ from population $j$. That is, if individual $p$ were the only individual corresponding to a particular ancestor then removing it would also remove the ancestor, thus necessitating these updates. The second term of Equation S.12 corresponds to allocating a new ancestor for individual $p$. By applying the appropriate scaling, the probability of allocating a new ancestor can be written as $\left(1 + \frac{(\gamma+m_{\cdot\cdot})(n'_{j\cdot k_{jp}} - m'_{\cdot k_{jp}}\sigma_j)}{(\theta_j + m'_{j\cdot}\sigma_j)(m'_{\cdot k_{jp}} - \alpha)}\right)^{-1}$. The procedure for updating each ancestral state is more precisely defined in Algorithm 1 in the main manuscript.

#### S1.4 Slice sampler

**Algorithm S1:** Slice sampler for concave log-probability function $l(x)$

```
Initialize x^(0), b_lower, b_upper
for i ← 1 to I do
    y ← l(x^(i−1)) ;
    b_l ← b_lower ;
    b_u ← b_upper ;
    Generate u ~ Unif(0, 1) ;
    y ← y + log(u) ;
    rejected ← TRUE ;
    while rejected do
        v ← Unif(0, 1) ;
        x^(try) ← b_l + v(b_l − b_u) ;
        if l(x^(try)) > y then
            x^(i) ← x^(try) ;
            rejected ← FALSE ;
        else
            if x^(try) < x^(i−1) then
                b_l ← x^(try) ;
            else
                b_u ← x^(try) ;
            end
        end
    end
end
```

### S2 Diversity Indices for the HPY Model

#### S2.1 Alpha-diversity

One of the virtues of the HPY process is that one can compute the Hill numbers Hill (1973) for integers greater than equal to 2 for the $j^{\text{th}}$ population,

$$^{q}Dj = \left(\sum_{i=1}^{\infty} p_i^q\right)^{\frac{1}{1-q}},$$

where $q > 1$ and $p_i$ is the relative abundance of the $i^{\text{th}}$ species; $1 - 1/(^2Dj)$ is Simpson's index (using the definition given in Equation 4 in the main manuscript), whereas with increasing $q$, greater weight is placed upon more abundant species. The sum $\sum_{i=1}^{\infty} p_i^q$ is the probability of sampling $q$ individuals of the same species when sampling with replacement, which, given estimates for the parameters $\alpha$, $\gamma$, $\theta_j$, and $\sigma_j$, can be done via the so-called "Chinese restaurant franchise" Teh (2006). We will consider two approaches: the first yields simple expressions by only taking into account the fitted parameters. The second, following Cerquetti (2015), offers a more informed estimate, also taking into account the samples used to obtain the parameter estimates. The latter yields considerably more complicated expressions, so we will only compute Simpson's index.

##### S2.1.1 Parameter-based estimates

First, consider Simpson's index. Suppose we have sampled one individual; there are two ways we could sample a second individual of the same species, either we sample a descendant of the first individual, with probability $\frac{1-\sigma_j}{\theta_j+1}$, or we sample a new ancestor of the same species from the metacommunity, with probability $\frac{\theta_j+\sigma_j}{\theta_j+1}\frac{1-\alpha}{\gamma+1}$. These are disjoint events, so their sum gives us the probability of sampling two distinct individuals, which gives us:

$$\frac{1}{^2Dj} = \frac{1-\sigma_j}{\theta_j+1} + \frac{\theta_j+\sigma_j}{\theta_j+1}\frac{1-\alpha}{\gamma+1}. \tag{S.13}$$

Now consider $q = 3$: the reasoning above gives us the probability that the second sampled individual is of the same species as the first. To obtain the probability that the third has the same type, we must now consider five scenarios. For the first two, consider the case when the second species was a descendant of the first; to be of the same species, the third can either be a descendent of the first or second individual, with probability $\frac{2-\sigma_j}{\theta_j+2}$, or they again can be a new ancestor of the same species from the metacommunity, with probability $\frac{\theta_j+\sigma_j}{\theta_j+2}\frac{1-\alpha}{\gamma+1}$. If, on the other hand, the first two individuals came from distinct ancestral lineages drawn from the metacommunity, then we must consider three possibilities: that the third is a descendent of the first ancestor, or of the second ancestor, both of which have probability $\frac{1-\sigma_j}{\theta_j+2}$, or the third is again a new ancestor of the same species drawn from the metacommunity with probability $\frac{\theta_j+2\sigma_j}{\theta_j+2}\frac{2-\alpha}{\gamma+2}$. The probability of drawing three individuals of the same type is thus

$$\frac{1-\sigma_j}{\theta_j+1}\left(\frac{2-\sigma_j}{\theta_j+2}+\frac{\theta_j+\sigma_j}{\theta_j+2}\frac{1-\alpha}{\gamma+1}\right)+\frac{\theta_j+\sigma_j}{\theta_j+1}\frac{1-\alpha}{\gamma+1}\left(2\frac{1-\sigma_j}{\theta_j+2}+\frac{\theta_j+2\sigma_j}{\theta_j+2}\frac{2-\alpha}{\gamma+2}\right)$$
$$=\frac{1-\sigma_j}{\theta_j+1}\frac{2-\sigma_j}{\theta_j+2}+3\frac{\theta_j+\sigma_j}{\theta_j+1}\frac{1-\sigma_j}{\theta_j+2}\frac{1-\alpha}{\gamma+1}+\frac{\theta_j+\sigma_j}{\theta_j+1}\frac{\theta_j+2\sigma_j}{\theta_j+2}\frac{1-\alpha}{\gamma+1}\frac{2-\alpha}{\gamma+2}.$$

Taking the reciprocal square root of the above yields ${}^3Dj$. Considering the latter expression allows us to quickly pass to the general case: each of the terms corresponds to the number of ancestors, one, two or three, and the probability that a sample is derived from $m$ ancestors is independent of the order in which we draw ancestors and descendants. We need only take into account the number of ways the individuals in the sample are partitioned into groups of descendants of those ancestors: here, there is only one way that all the individuals can have all the same ancestor, or all distinct ancestors, while there are $\binom{3}{2} = 3$ ways that two of the three individuals share a common ancestor while the third starts its own ancestral line.

Recall that the number of ways that $q$ individuals can be partitioned into $m$ labelled subsets (grouped by ancestor) of size $q_1, \ldots, q_m$ is the multinomial coefficient

$$\binom{q}{q_1, \ldots, q_m} = \frac{q!}{q_1! \cdots q_m!}.$$

This, however, multiply counts partitions with more than one subset of the same size (*e.g.* $\frac{3!}{1!1!1!} = 6$, because $\{\{1\},\{2\},\{3\}\}$, $\{\{2\},\{1\},\{3\}\}$, *etc.* are counted as distinct partitions. To that end, we instead set $a_r$ to be the number of parts of subsets of size $r$, $r = 1, \ldots, q$, so that $a_1 + \cdots + a_q = m$ and $a_1 + 2a_2 + \cdots + qa_q = q$. We then have that the number of unlabelled subsets is

$$\frac{q!}{\prod_{r=1}^{q}(r!)^{a_r} a_r!}.$$

We can quickly derive an expression for ${}^qDj$. The probability of drawing $m$ ancestors of the same species from the metacommunity is

$$\frac{1-\alpha}{\gamma+1} \cdots \frac{m-1-\alpha}{\gamma+m-1},$$

whereas the probability that $q$ individuals are descended from the same ancestor, $q = 1, \ldots, m$, and $m$ ancestors are derived from the metacommunity is (independently of the order of these events)

$$\frac{(\theta_j+\sigma_j)\cdots(\theta_j+(m-1)\sigma_j)\prod_{i=1}^{m}(1-\sigma_j)\cdots(q_i-1-\sigma_j)}{(\theta_j+1)\cdots(\theta_j+q-1)}.$$

Given $m$, let $A_{m,q} = \{\{a_1, \ldots, a_q\} : a_1 + \cdots + a_q = m, a_1 + 2a_2 + \cdots + qa_q = q\}$. Summing over all possible numbers of ancestors and partitions, and the number of ways of grouping individuals into those partitions yields (for $q > 1$):

$${}^qDj = \left( \sum_{m=1}^{q} \sum_{\{a_1,\ldots,a_q \in A_{m,q}\}} \frac{q!}{\prod_{r=1}^{q}(r!)^{a_r} a_r!} \frac{(1-\alpha)^{[m-1]}}{(1+\gamma)^{[m-1]}} \prod_{r=1}^{q} \left[(1-\sigma_j)^{[r-1]}\right]^{a_r} \frac{\prod_{k=0}^{m-1}(\theta_j+k\sigma_j)}{\theta_j^{[q]}} \right)^{\frac{1}{1-q}},$$

where $x^{[n]} = x(x+1)\cdots(x+n-1)$ is the rising factorial and where we use the conventions that $x^{[0]} = 1$ and $\prod_{k=1}^{0} f(x_k) = 1$.

##### S2.1.2 Parameter and sample based estimates

Now suppose we have already sampled $n_{jtk}$ descendants, $t = 1,\ldots,m_{jk}$ of $m_{jk}$ ancestors of species $k$, $k = 1,\ldots,K$ in the $j^{\text{th}}$ community, $j = 1\ldots,J$. Proceeding as above we can then obtain a form for Simpson's index conditional on this additional information: now, we ask the probability that the next two sampled individuals are of the same species. Our sampling across multiple local communities has given us additional information about the frequencies of species both in the community of interest, $j$, and also of species abundances in the metacommunity. As a consequence, we can no longer consider the first of the two individuals sampled *ex nihilo*, but must instead consider two scenarios: either they are a descendent of one of the $m_{j\cdot}$ ancestors in community $j$, say $t$, with probability $\frac{n_{j,t}-\sigma_j}{\theta+n_{j\cdot}}$, or they are drawn from the metacommunity with probability $\frac{\theta_j+m_{j,\cdot}\sigma_j}{\theta+n_{j\cdot}}$. In the latter case, we must consider two possibilities: that they are of a species, say $k$, of the $K$ previously observed in one one of the local communities, with probability $\frac{m_{\cdot k}-\alpha}{\gamma+m_{\cdot\cdot}}$, or that they are of a novel species, previously unobserved, with probability $\frac{\gamma+K\alpha}{\gamma+m_{\cdot\cdot}}$.

Again, we consider the possible scenarios in which the second individual is of the same species, which in turn depend on the identity of the first. If the first was of a novel species, then to be of the same species the second must either be a descendent of the first, with probability $\frac{1-\sigma_j}{\theta+n_{j\cdot}+1}$, or drawn from the metacommunity and of the same new species, with probability $\frac{\theta_j+(m_{j,\cdot}+1)\sigma_j}{\theta+n_{j\cdot}+1}\frac{1-\alpha}{\gamma+m_{\cdot\cdot}+1}$. If the first was of a previously sampled species, there are three possibilities for the second to be of the same species: they could again be a descendant of the first, with probability $\frac{1-\sigma_j}{\theta+n_{j\cdot}+1}$, they could be the descendent of a different ancestor $t'$ from the first of the same species, with probability $\frac{n_{j,t'}-\sigma_j}{\theta+n_{j\cdot}+1}$, or they could be a new ancestor of the same species drawn from the metacommunity, with probability $\frac{\theta_j+m_{j,\cdot}\sigma_j}{\theta+n_{j\cdot}+1}\frac{m_{\cdot k}-\alpha}{\gamma+m_{\cdot\cdot}}$ if the first was descended from a previously sampled ancestor, and probability $\frac{\theta_j+(m_{j,\cdot}+1)\sigma_j}{\theta+n_{j\cdot}+1}\frac{m_{\cdot k}+1-\alpha}{\gamma+m_{\cdot\cdot}+1}$

if the first was also a new ancestor sampled from the metacommunity.

Summing these probabilities yields expression for Simpson's index in population $j$ conditional on the previous samples:

$$
\begin{aligned}
\frac{1}{{}^2Dj} = \sum_{t=1}^{m_{j\cdot}} \Bigg\{ & \frac{n_{jt}-\sigma_j}{\theta_j+n_{j\cdot}} \left( \frac{(n_{jt}+1)-\sigma_j}{\theta_j+n_{j\cdot}+1} + \sum_{t':\psi_{jt'}=\psi_{jt},t'\neq t} \frac{n_{jt'}-\sigma_j}{\theta_j+n_{j\cdot}+1} + \frac{\theta_j+m_{j\cdot}\sigma_j}{\theta_j+n_{j\cdot}+1}\frac{m_{\cdot\psi_{jt}}-\alpha}{\gamma+m_{\cdot\cdot}} \right) \Bigg\} \\
&+ \frac{\theta_j+m_{j\cdot}\sigma_j}{\theta_j+n_{j\cdot}} \Bigg( \frac{\gamma+K\alpha}{\gamma+m_{\cdot\cdot}} \left[ \frac{1-\sigma_j}{\theta_j+n_{j\cdot}+1} + \frac{\theta_j+(m_{j\cdot}+1)\sigma_j}{\theta_j+n_{j\cdot}+1}\frac{1-\alpha}{\gamma+m_{\cdot\cdot}+1} \right] \\
&+ \sum_{k=1}^{K} \left\{ \frac{m_{\cdot k}-\alpha}{\gamma+m_{\cdot\cdot}} \left[ \frac{1-\sigma_j}{\theta_j+n_{j\cdot}+1} + \sum_{t':\psi_{jt'}=k} \frac{n_{jt'}-\sigma_j}{\theta_j+n_{j\cdot}+1} + \frac{\theta_j+(m_{j\cdot}+1)\sigma_j}{\theta_j+n_{j\cdot}+1}\frac{(m_{\cdot k}+1)-\alpha}{\gamma+m_{\cdot\cdot}+1} \right] \right\} \Bigg) \\
= & \frac{\sum_{t=1}^{m_{j\cdot}}(n_{jt}-\sigma_j)[n_{j\cdot\psi_{jt}}-(m_{j\psi_{jt}}+1)\sigma_j]}{(\theta_j+n_{j\cdot})(\theta_j+n_{j\cdot}+1)} \\
&+ (\theta_j+m_{j\cdot}\sigma_j) \Bigg\{ \frac{\sum_{t=1}^{m_{j\cdot}}(n_{jt}-\sigma_j)(m_{\cdot\psi_{jt}}-\alpha)}{(\theta_j+n_{j\cdot})(\theta_j+n_{j\cdot}+1)(\gamma+m_{\cdot\cdot})} \\
&\quad + \frac{(\gamma+K\alpha)(1-\sigma_j)+\sum_{k=1}^{K}(m_{\cdot k}-\alpha)[n_{j\cdot k}+1-(m_{jk}+1)\sigma_j]}{(\theta_j+n_{j\cdot})(\theta_j+n_{j\cdot}+1)(\gamma+m_{\cdot\cdot})} \\
&\quad + \frac{(\gamma+K\alpha)[\theta_j+(m_{j\cdot}+1)\sigma_j](1-\alpha)+\sum_{k=1}^{K}(m_{\cdot k}-\alpha)[\theta_j+(m_{j\cdot}+1)\sigma_j](m_{\cdot k}+1-\alpha)}{(\theta_j+n_{j\cdot})(\theta_j+n_{j\cdot}+1)(\gamma+m_{\cdot\cdot})(\gamma+m_{\cdot\cdot}+1)} \Bigg\}.
\end{aligned}
\tag{S.14}
$$

The latter equality in Equation S.14 reflects a more computationally efficient way to calculate $1/({}^2Dj)$, as the first definition contains several unnecessary sums. One could in principle compute the conditional Hill numbers for integers $q > 2$ in the same manner; we will not.

#### S2.2 Beta-diversity

Likewise, we can consider Simpson's index as a measure of beta-diversity, *i.e.* comparing species diversity across populations. In this case we wish to calculate the probability of sampling one species in population $j$, then subsequently sampling the same species in population $j'$. Using only the parameters, and not previous samples, this is the probability that the second individual is drawn the same species in the metacommunity,

$$\frac{1}{{}^2Djj'} = \frac{1-\alpha}{\gamma+1}. \tag{S.15}$$

If we condition on previous samples, we need to take into account that previously existing ancestors of the same species in communities $j$ and $j'$. Proceeding as for our calculations for the conditional alpha-diversity above, we obtain

$$\begin{aligned}
\frac{1}{{}^2Djj'} &= \sum_{t=1}^{m_{j\cdot}} \left\{ \frac{n_{jt}-\sigma_j}{\theta_j+n_{j\cdot}} \left( \sum_{t':\psi_{j't'}=\psi_{jt}} \frac{n_{j't'}-\sigma_{j'}}{\theta_{j'}+n_{j'\cdot}} + \frac{\theta_{j'}+m_{j'\cdot}\sigma_{j'}}{\theta_{j'}+n_{j'\cdot}} \frac{m_{\cdot\psi_{jt}}-\alpha}{\gamma+m_{\cdot\cdot}} \right) \right\} \\
&\quad + \frac{\theta_j+m_{j\cdot}\sigma_j}{\theta_j+n_{j\cdot}} \left( \frac{\gamma+K\alpha}{\gamma+m_{\cdot\cdot}} \left[ \frac{\theta_{j'}+m_{j'\cdot}\sigma_{j'}}{\theta_{j'}+n_{j'\cdot}} \frac{1-\alpha}{\gamma+m_{\cdot\cdot}+1} \right] \right. \\
&\quad \left. + \sum_{k=1}^{K} \frac{m_{\cdot k}-\alpha}{\gamma+m_{\cdot\cdot}} \left[ \sum_{t:\psi_{j't}=k} \frac{n_{j't}-\sigma_{j'}}{\theta_{j'}+n_{j'\cdot}} + \frac{\theta_{j'}+m_{j'\cdot}\sigma_{j'}}{\theta_{j'}+n_{j'\cdot}} \frac{m_{\cdot k}+1-\alpha}{\gamma+m_{\cdot\cdot}+1} \right] \right) \\
&= \frac{\sum_{t=1}^{m_{j\cdot}} (n_{jt}-\sigma_j)(n_{j'\cdot\psi_{jt}} - m_{j'\psi_{jt}}\sigma_{j'})}{(\theta_j+n_{j\cdot})(\theta_{j'}+n_{j'\cdot})} + (\theta_{j'}+m_{j'\cdot}\sigma_{j'}) \frac{\sum_{t=1}^{m_{j\cdot}} (n_{jt}-\sigma_j)(m_{\cdot\psi_{jt}}-\alpha)}{(\theta_j+n_{j\cdot})(\theta_{j'}+n_{j'\cdot})(\gamma+m_{\cdot\cdot})} \\
&\quad + (\theta_j+m_{j\cdot}\sigma_j) \left\{ \frac{\sum_{k=1}^{K} (m_{\cdot k}-\alpha)(n_{j'\cdot k} - m_{j'k}\sigma_{j'})}{(\theta_j+n_{j\cdot})(\theta_{j'}+n_{j'\cdot})(\gamma+m_{\cdot\cdot})} \right. \\
&\quad \left. + \frac{(\gamma+K\alpha)(\theta_{j'}+m_{j'\cdot}\sigma_{j'})(1-\alpha) + (\theta_{j'}+m_{j'\cdot}\sigma_{j'}) \sum_{k=1}^{K} (m_{\cdot k}-\alpha)(m_{\cdot k}+1-\alpha)}{(\theta_j+n_{j\cdot})(\theta_{j'}+n_{j'\cdot})(\gamma+m_{\cdot\cdot})(\gamma+m_{\cdot\cdot}+1)} \right\}.
\end{aligned} \tag{S.16}$$

Once again, the latter expression in Equation S.16 is a more computationally efficient expression. Thus, if we have a species abundance table over multiple populations, we could run the HPY Gibbs sampler to obtain samples of the HPY parameters, and use the above expression to obtain estimates for each of Simpson's alpha-diversity and beta-diversity indices.

### S3 Supplementary figures for simulation

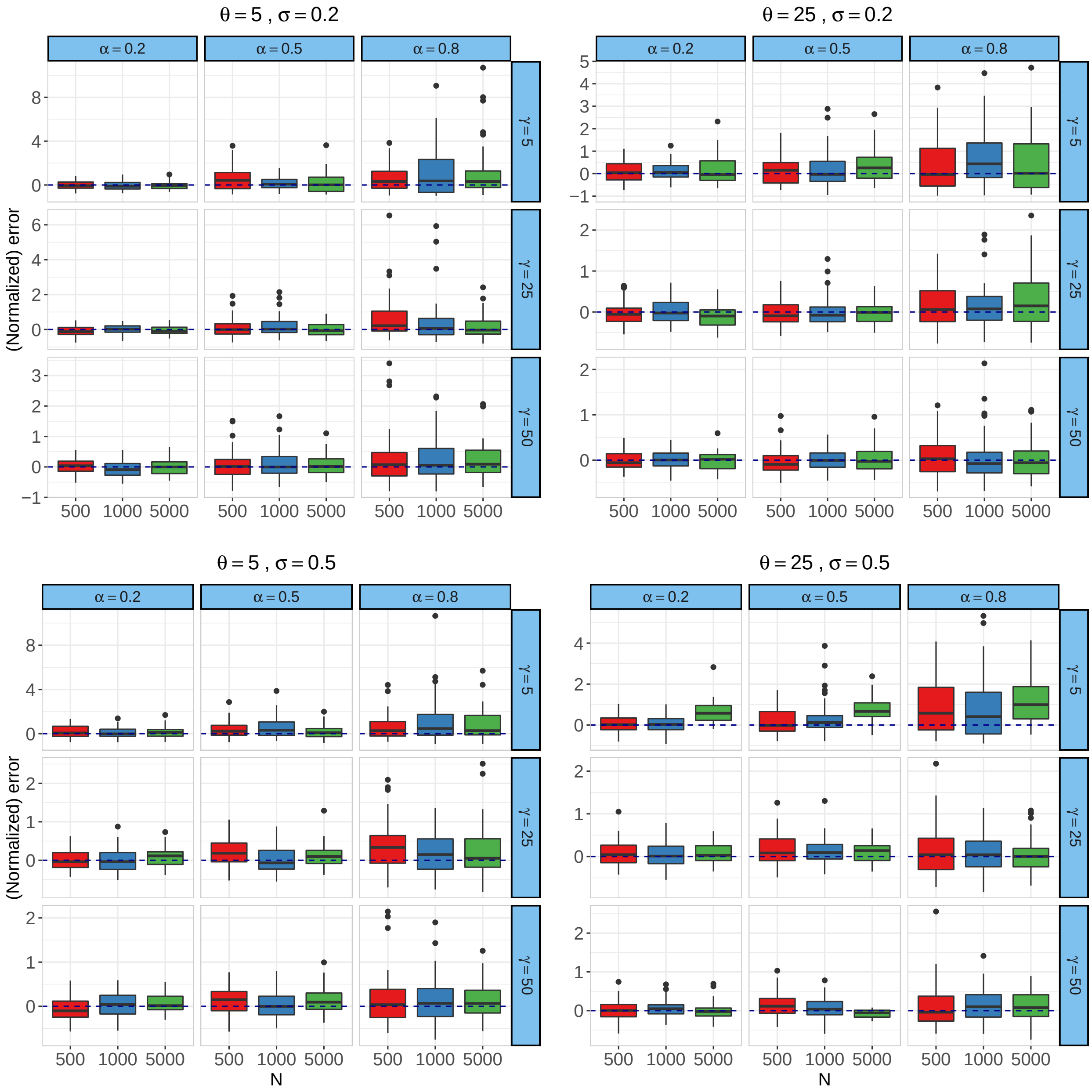


Figure S1: Normalized estimation error for top-level concentration ($\gamma$) in the simulation scenarios.

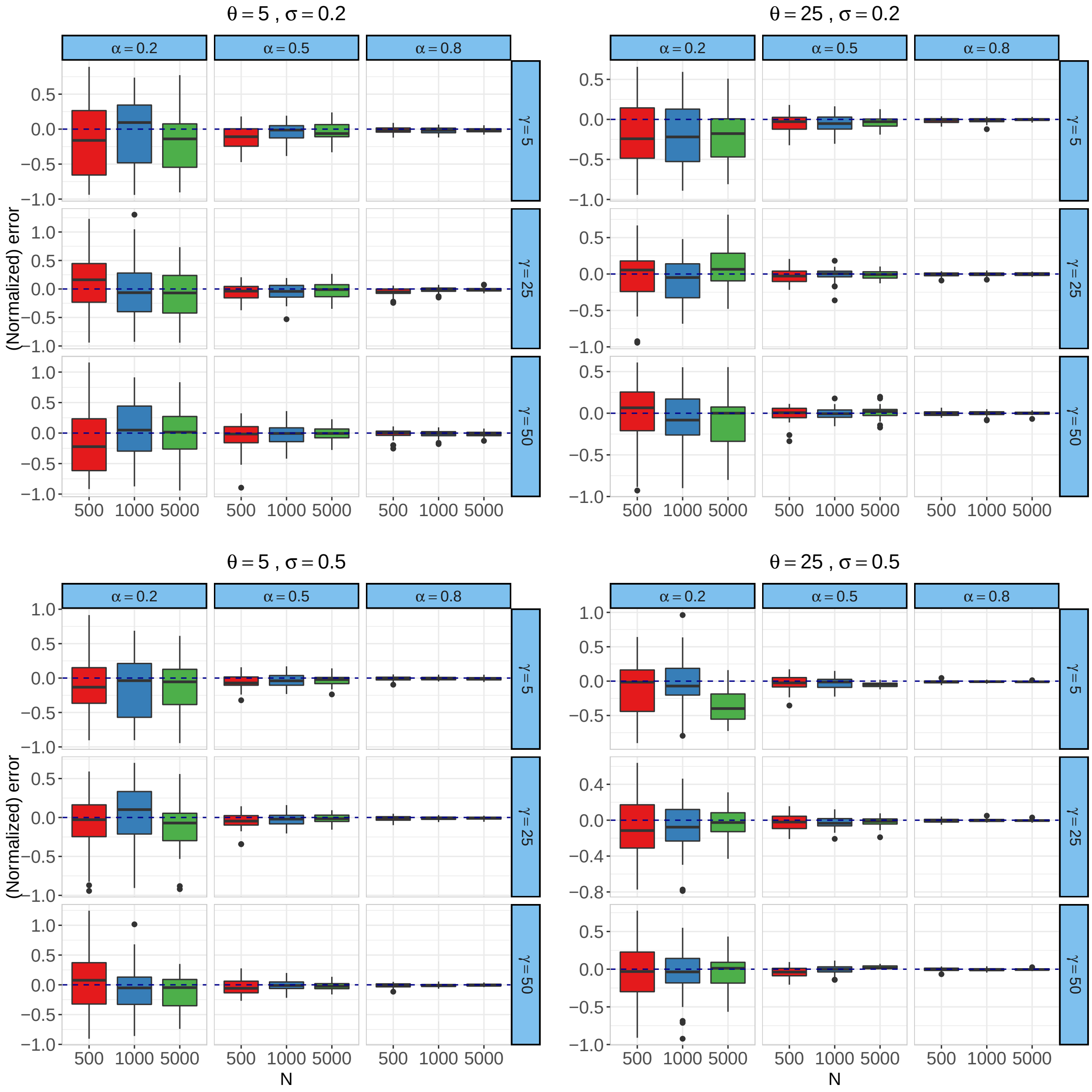


Figure S2: Normalized estimation error for top-level discount ($\alpha$) in the simulation scenarios.

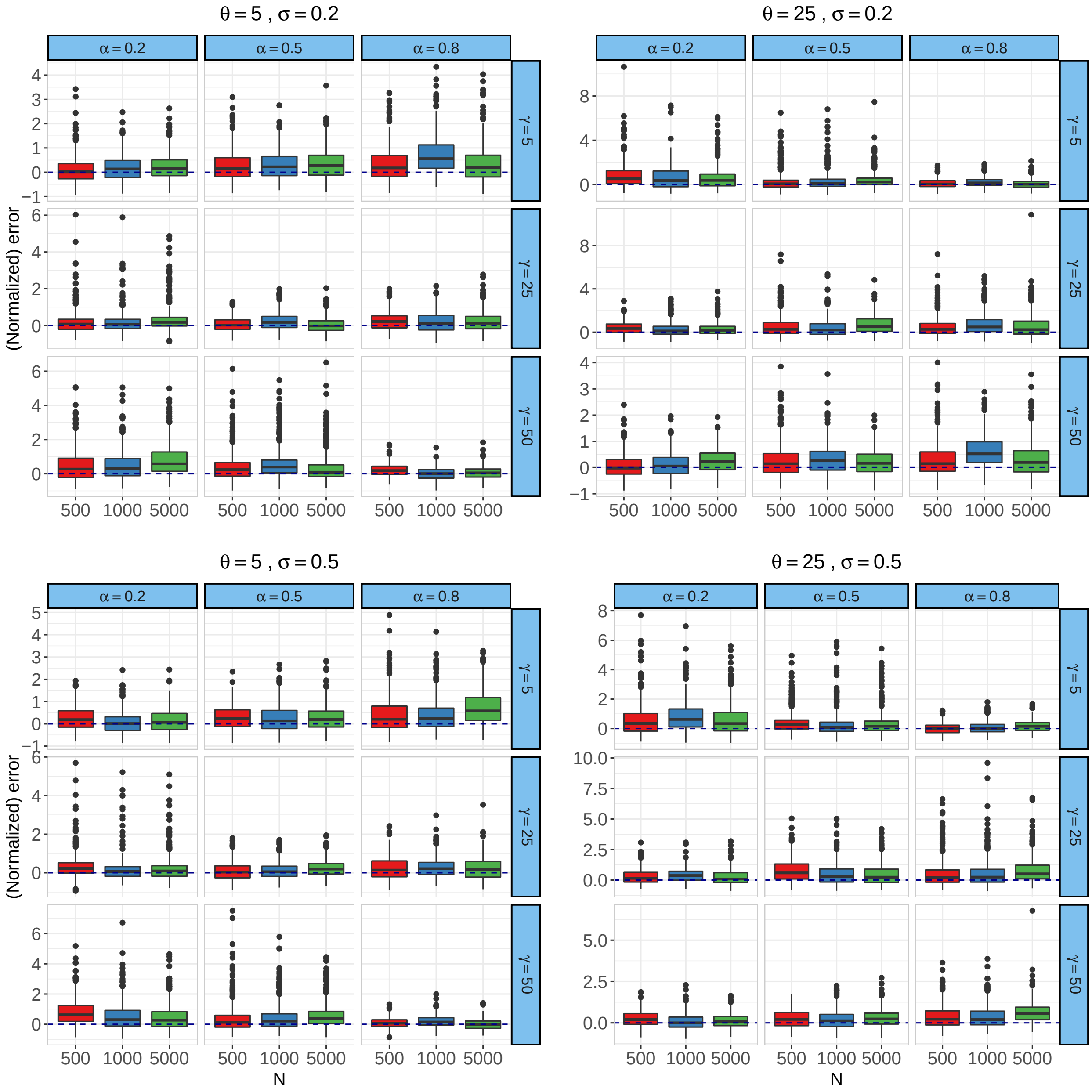


Figure S3: Normalized estimation error for local-level concentrations ($\theta_j$) in the simulation scenarios.

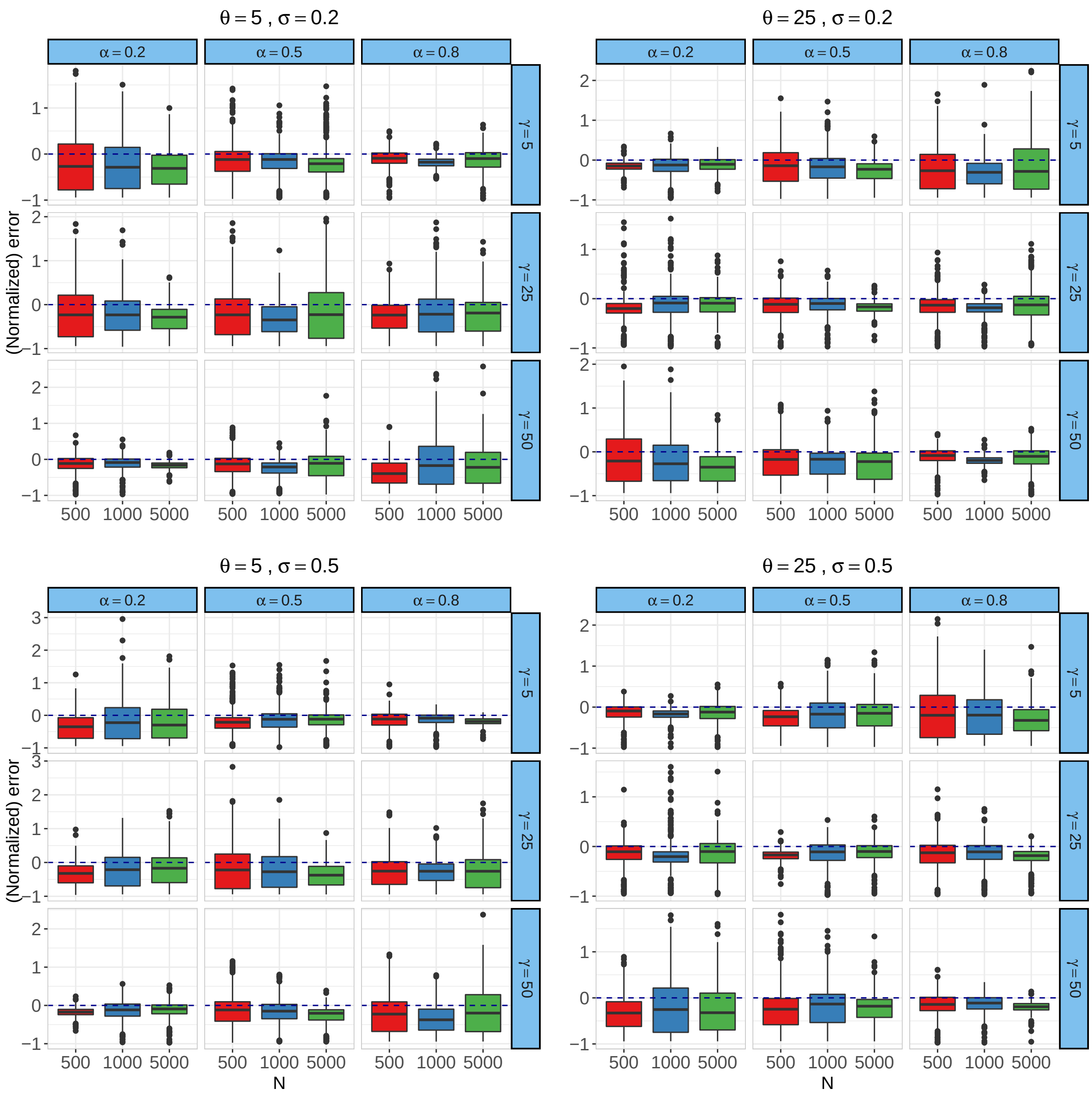


Figure S4: Normalized estimation error for local-level discounts ($\sigma_j$) in the simulation scenarios.

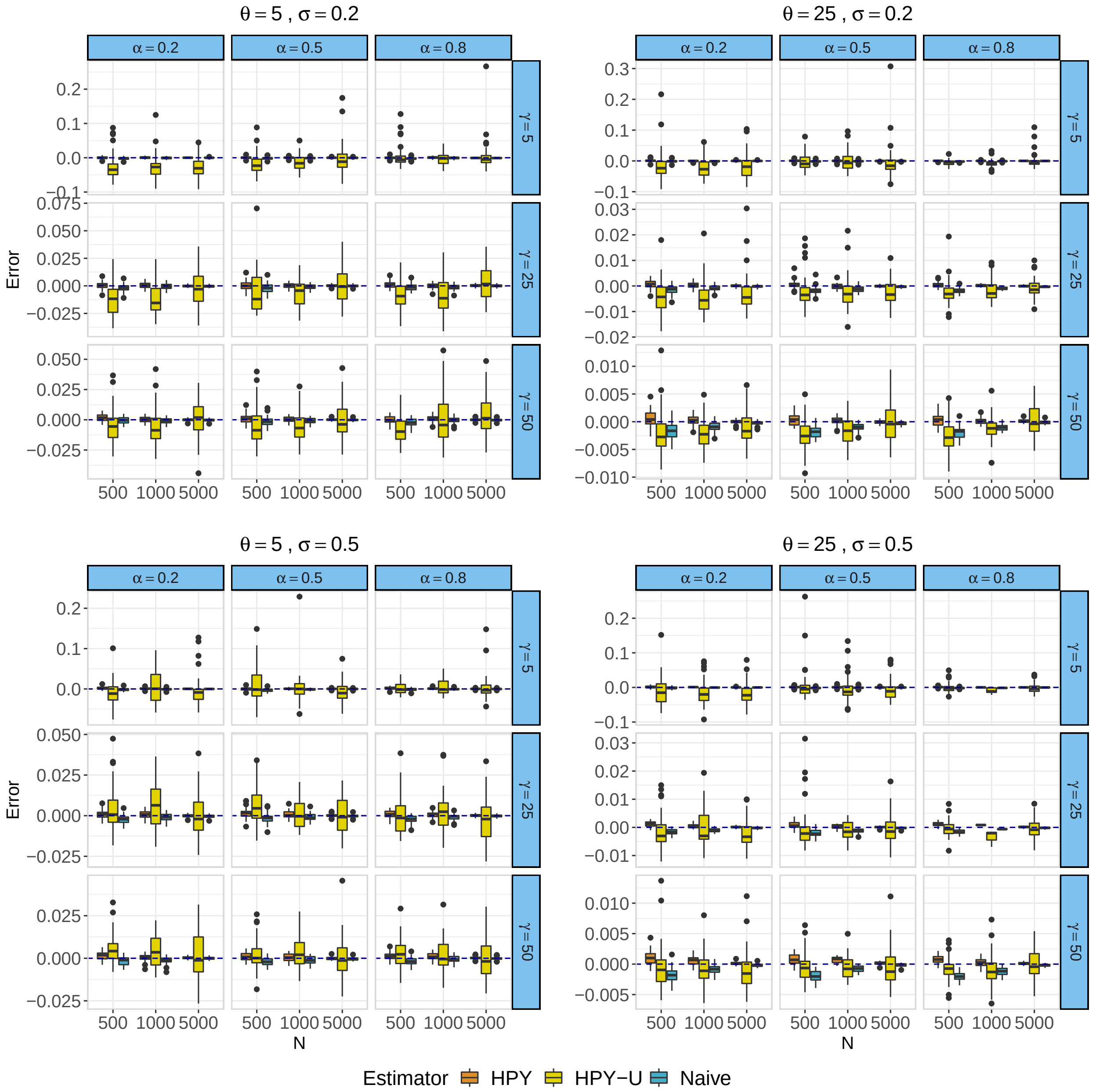


Figure S5: Estimation error for Simpson's index (alpha-diversity) in the simulation scenarios comparing conditional Simpson's index (HPY), unconditional Simpson's index (HPY-U) and naïve estimates.

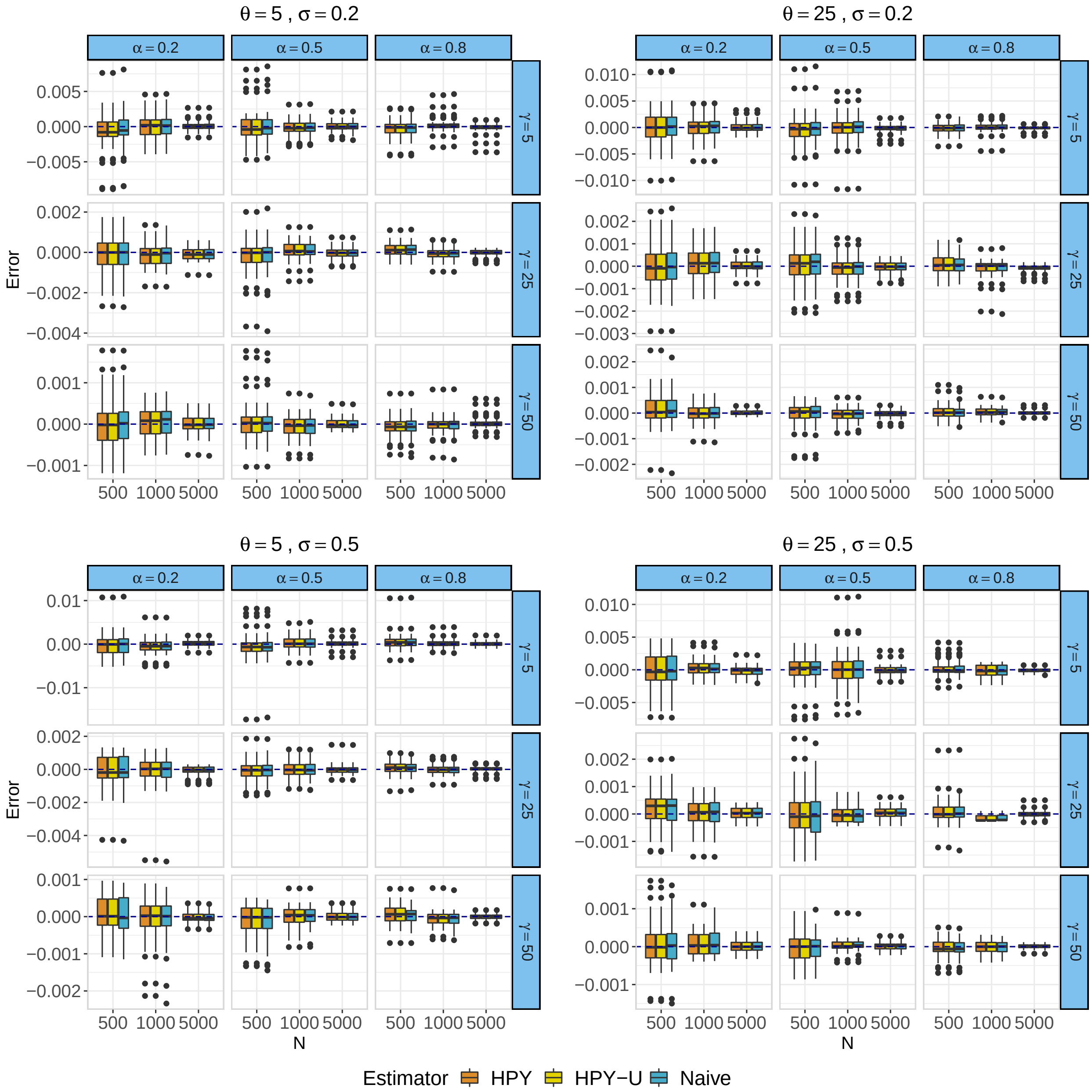


Figure S6: Estimation error for Simpson's index (beta-diversity) in the simulation scenarios

### S4 Supplementary figures for data analysis

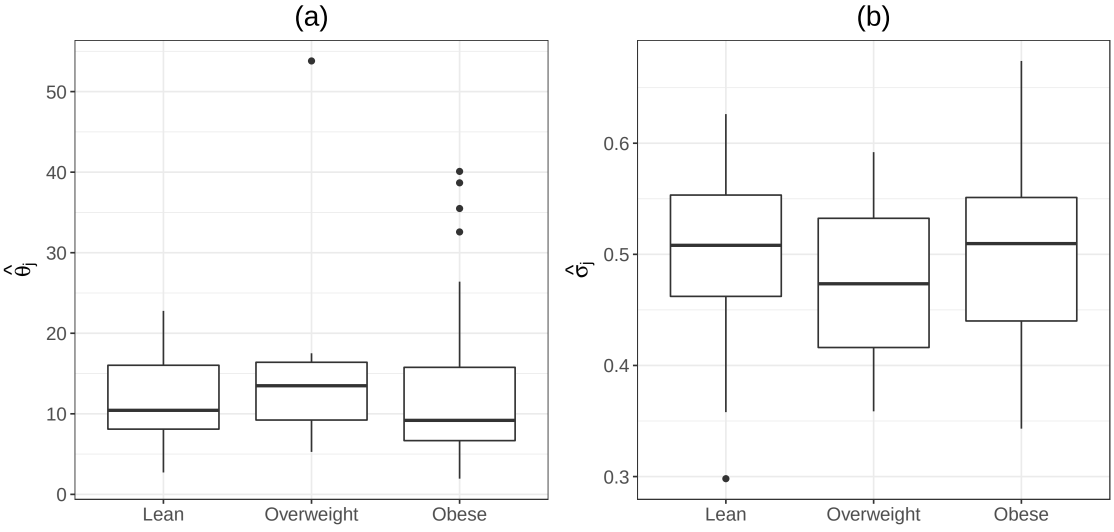


Figure S7: **(a)** Estimated local concentration parameter ($\widehat{\theta}_j$) in the obesity categories. **(b)** Estimated local concentration parameter ($\widehat{\sigma}_j$) in the obesity categories.

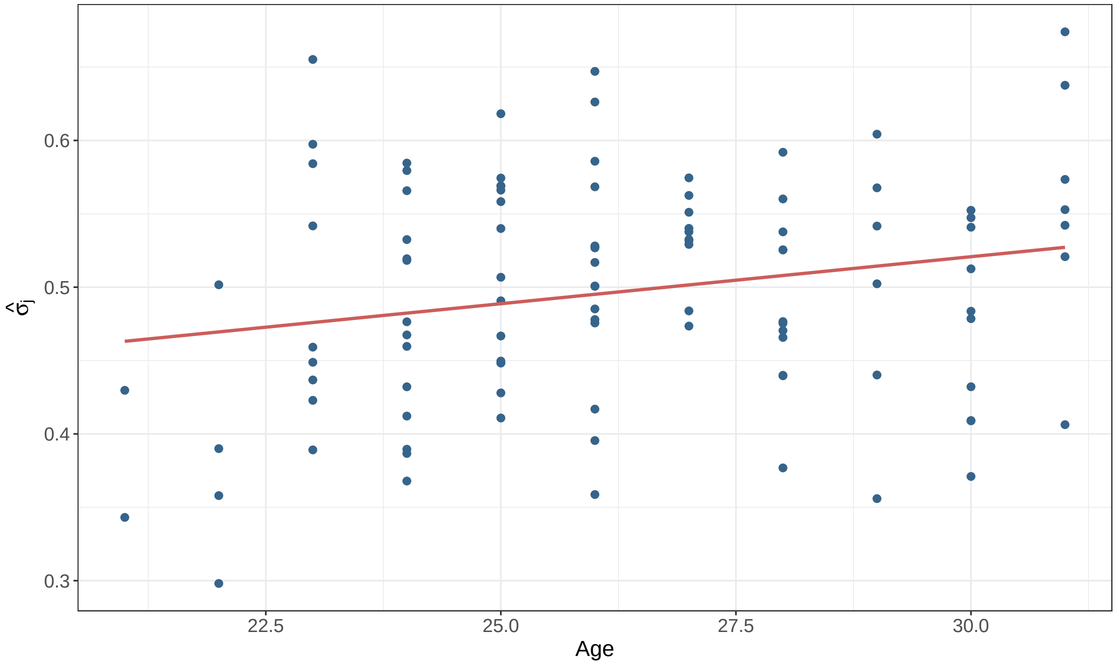


Figure S8: Estimated local discount parameter ($\widehat{\sigma}_j$) vs. age in the lean/obese twin study.

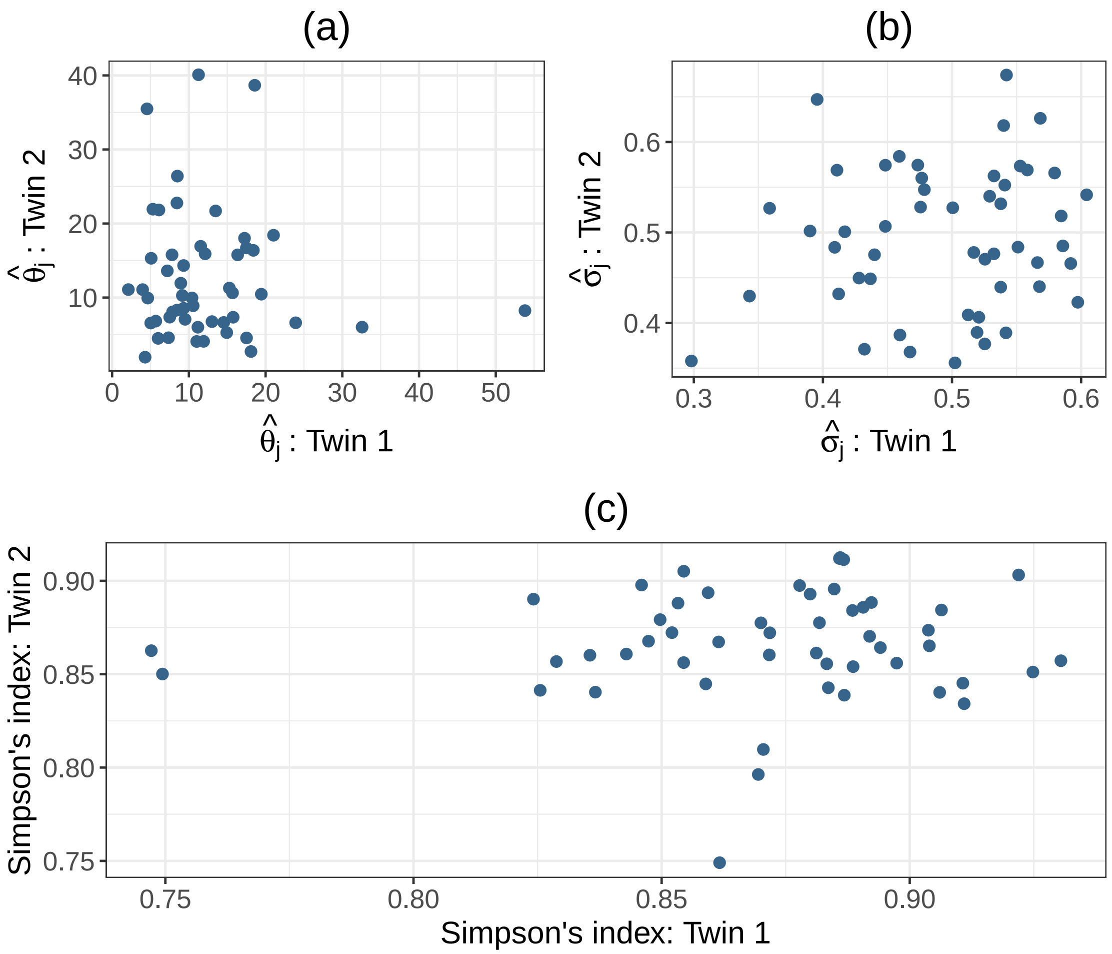


Figure S9: Concordance between twin pairs for: **(a)** Local concentration ($\widehat{\theta}_j$), **(b)** Local discount ($\widehat{\sigma}_j$), and **(c)** Simpson's index

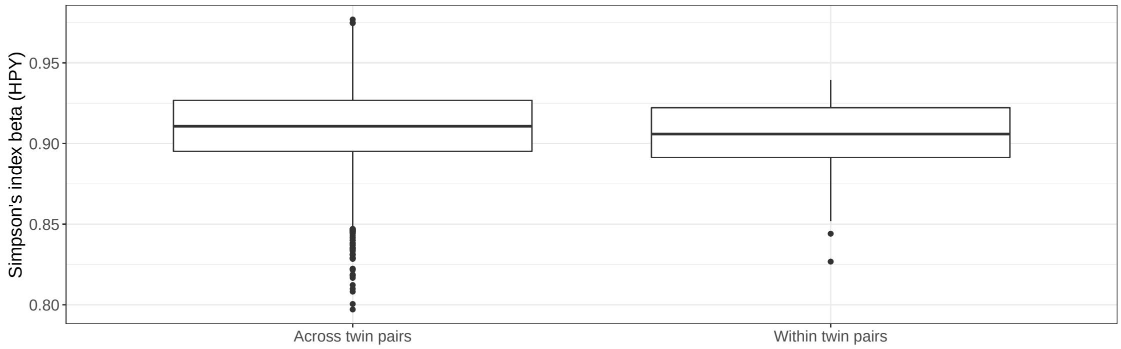


Figure S10: Comparing Simpson's index beta calculated between twin pairs and within twin pairs.

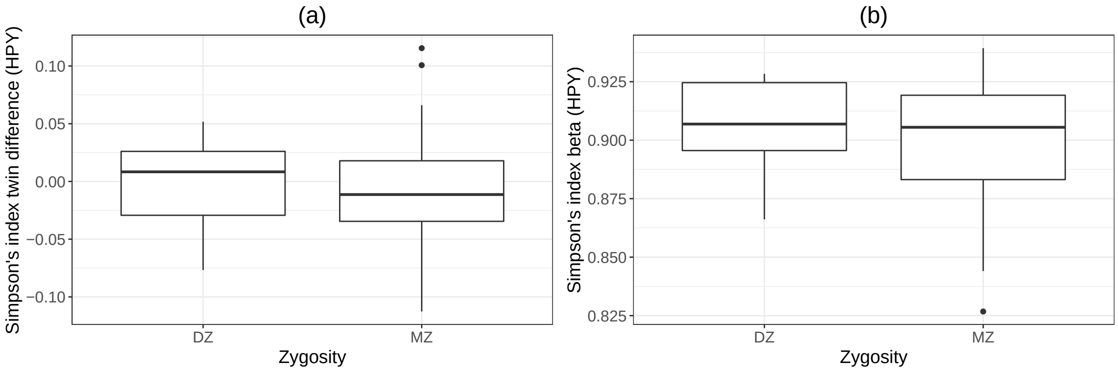


Figure S11: Comparing **(a)** Simpson's index differences between monozygotic (MZ) and dizygotic (DZ) twin pairs; and **(b)** Simpson's index beta within MZ and DZ pairs.

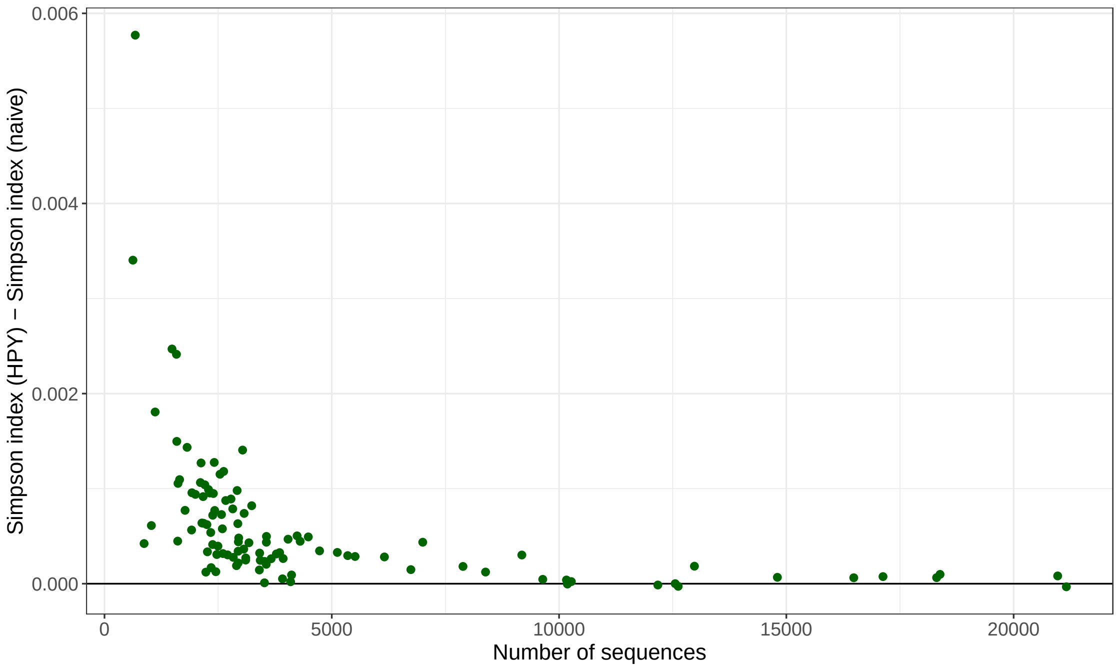


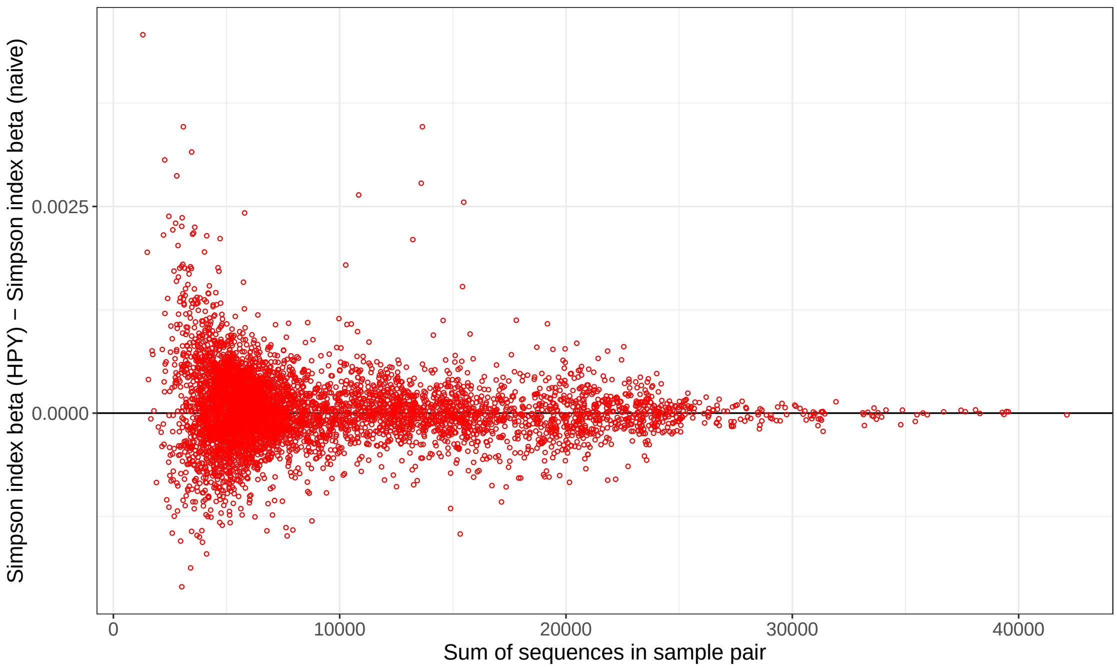


Figure S12: Difference between Simpson's index for **(top)** alpha-diversity and **(bottom)** beta-diversity calculated using HPY model and naïvely as a function of the number of sequences in each sample in the obese/lean twins study. For the beta-diversity plot, the sum of the number of sequences for each pair of samples is considered.

Wray Buntine and Marcus Hutter. A bayesian view of the poisson-dirichlet process. *arXiv preprint arXiv:1007.0296*, 2010. 2

Wray Buntine. Hyper-parameter estimation for the dirichlet prior. 2012. 2, 3, 4

Michael D Escobar and Mike West. Bayesian density estimation and inference using mixtures. *Journal of the american statistical association*, 90(430):577–588, 1995. 2

Marco Battiston, Stefano Favaro, and Yee Whye Teh. Multi-armed bandit for species discovery: a bayesian nonparametric approach. *Journal of the American Statistical Association*, 113(521):455–466, 2018. 5

Mark O Hill. Diversity and evenness: a unifying notation and its consequences. *Ecology*, 54 (2):427–432, 1973. 7

Yee Whye Teh. A hierarchical bayesian language model based on pitman-yor processes. In *Proceedings of the 21st International Conference on Computational Linguistics and 44th Annual Meeting of the Association for Computational Linguistics*, pages 985–992, 2006. 7

Annalisa Cerquetti. Bayesian estimation of gini-simpson's index under mainland-island community structure. In *Convegno della Società Italiana di Statistica*, pages 61–70. Springer, 2015. 7
